# Mature leaves produce a multi-layered wound periderm by integrating phytohormone signaling with ATML1-mediated epidermal specification

**DOI:** 10.1101/2024.09.02.607870

**Authors:** Jung-Min Lee, Woo-Taek Jeon, Minsoo Han, Myung Kwon, Kyungyoon Kim, Sujeong Je, Hoon Jung, Geon Heo, Yasuyo Yamaoka, Yuree Lee

## Abstract

The epidermis of plants forms a protective barrier against various stress, but how breaches in the epidermis are repaired is not well understood. Here, we investigated wound healing in the mature leaves of *Arabidopsis*. We discover a novel type of wound periderm comprising a multi-layered ligno-suberized barrier covered with cuticular wax, which is formed by mesophyll cells that adopt an epidermal fate. Mesophyll cells of protective layer 1 (P1), just beneath the wound, transition into epidermal cells, which seal the wound by depositing cuticle. As P1 undergoes cell death, protective layer 2 (P2), which underlies P1, takes the place of P1 and undergoes ligno-suberization. This multi-layered periderm involves integration of ethylene and jasmonic acid signaling with ATML1, a key transcription factor in epidermal specification, to coordinate cell layer-specific functions. This novel wound periderm also occurs in the leaves of tobacco and *Capsella*, suggesting it is a widespread phenomenon.

## INTRODUCTION

Plants living a sessile existence in the soil endure various biological and environmental stresses throughout their lives, often facing mechanical injuries. Wounds can significantly harm plants by causing dehydration through solute leakage and heightening their susceptibility to pathogen infections. Therefore, timely and effective management of wounds is essential for plant survival.

While extensive research has explored the signaling networks involved in the wound response (Leon et al., 2001; Miller et al., 2009; Mousavi et al., 2013; Zhang et al., 2019), understanding their precise roles in local wound healing remains limited. Plants employ various mechanisms to seal wounds, including the deposition of impermeable substances, such as lignin and suberin (Barros et al., 2015; Moon et al., 1984; Serra and Geldner, 2022). The composition of these substances may vary depending on the species and organs involved (Rittinger et al., 1987; Savatin et al., 2014). For instance, in potato tubers and fruits, the wound healing process consists of forming a protective layer known as the wound periderm, which entails the deposition of lignin and suberin (Graca and Santos, 2006; Lulai et al., 2008; Tao et al., 2016). Similarly, in cherry foliage and carrot roots, wounds induce the development of a distinct ligno-suberized boundary layer, where the cell wall initially undergoes lignification, and then forms suberin lamellae (Espelie et al., 1980; Rittinger et al., 1987). Currently, research on wound healing concentrates mostly on tubers and fruits, with little investigation of the healing of leaf wounds.

The repair of wound-related tissue damage is sometimes accompanied by stem cell activation and regeneration, governed by various plant hormones (Christiaens et al., 2021; Ikeuchi et al., 2017; Lup et al., 2016). In explants of *Arabidopsis* leaves, wound-induced jasmonic acid (JA) triggers root organogenesis. Within two hours of leaf detachment, JA activates the ETHYLENE RESPONSE FACTOR109 (ERF109), which subsequently upregulates expression of *ANTHRANILATE SYNTHASE* α*1* (*ASA1*), a gene involved in tryptophan biosynthesis and auxin production. This JA–ERF109–*ASA1* wound signaling pathway enhances auxin biosynthesis, ultimately promoting root regeneration (Zhang et al., 2019). Single-cell ablation studies in roots have shed light on the cellular mechanisms involved in replenishing damaged cells. This process requires stem cell activation, regeneration, and expansion of neighboring cells, facilitated by localized increases in auxin accumulation and *ERF115* expression (Canher et al., 2020; Hoermayer et al., 2020; Zhou et al., 2019). In flowering stems, wound healing involves the transformation of differentiated cells into cambium (Matsuoka et al., 2021). These cambium-like cells proliferate and actively contribute to the tissue reunion process, regulated by transcription factors ANAC071 and its homolog ANAC096.

This study aimed to discover the cellular and molecular mechanisms that govern wound healing at sites of local physical injury in mature *Arabidopsis* leaves. Our findings reveal unexpected cuticle deposition during the wound-healing process. Also, the wound-healing process involves the coordinated action of multiple cell layers. The mesophyll cell layer directly beneath the damaged area, called protective layer 1 (P1), differentiates to form epidermal cells that form cuticle. Meanwhile, the underlying P2 undergoes ligno-suberization, forming another protective layer. Notably, cell death in P1 preceeds the expansion and ligno-suberization of P2, resulting in the ligno-suberized P2 layer becoming the outermost living layer covered by the cuticle. By employing genetic and pharmacological approaches, we elucidate the signaling components that govern spatiotemporal activation and cell death in these two layers. These findings offer a new perspective on the dynamics of wound healing and identify a novel type of wound periderm.

## RESULTS

### Wounding induces multilayered reorganization of mesophyll cells in mature leaves

To investigate the wound healing process in mature leaves, we made a precise cut on the outer edge of the eighth leaf of 4-week-old *Arabidopsis* plants and analysed the wounds by microscopy after staining with toluidine blue to assess permeability (Figure 1A). Quantification of the stained area revealed that wound healing began within the first day and progressed substantially over the following five days (Figure 1B). We used scanning electron microscopy to observe progression of the healing process at greater magnification. Immediately after wounding, mesophyll cells and internal airspace were visible in the leaf tissue. One day later, however, the cut surface was beginning to be filled in, and by the fifth day post-wounding (5 DPW), the surface was covered with a smooth, protective substance (Figure 1C). Three-dimensional quantitative analysis using micro-computed tomography revealed a significant loss of porosity in the region proximal to the wound (Figures 1D, 1E, and S1A), indicating that the sealing process involves more than simply cell wall remodeling in the surface layer. The airspace between mesophyll cells, the smooth surface, and loss of porosity at the wound site together suggested that the sealing process may involve complex changes, including cell division or expansion. Indeed, sections through the leaf adjacent to the wound revealed cellular rearrangement at the injured surface (Figure 1F). Under normal conditions, palisade mesophyll cells elongate in the adaxial–abaxial direction (from the upper to the lower side of the leaf). At the wound site, however, the mesophyll cells elongated in the transverse direction, toward the wound. No evident signs of cell division were observed.

**Figure 1.**
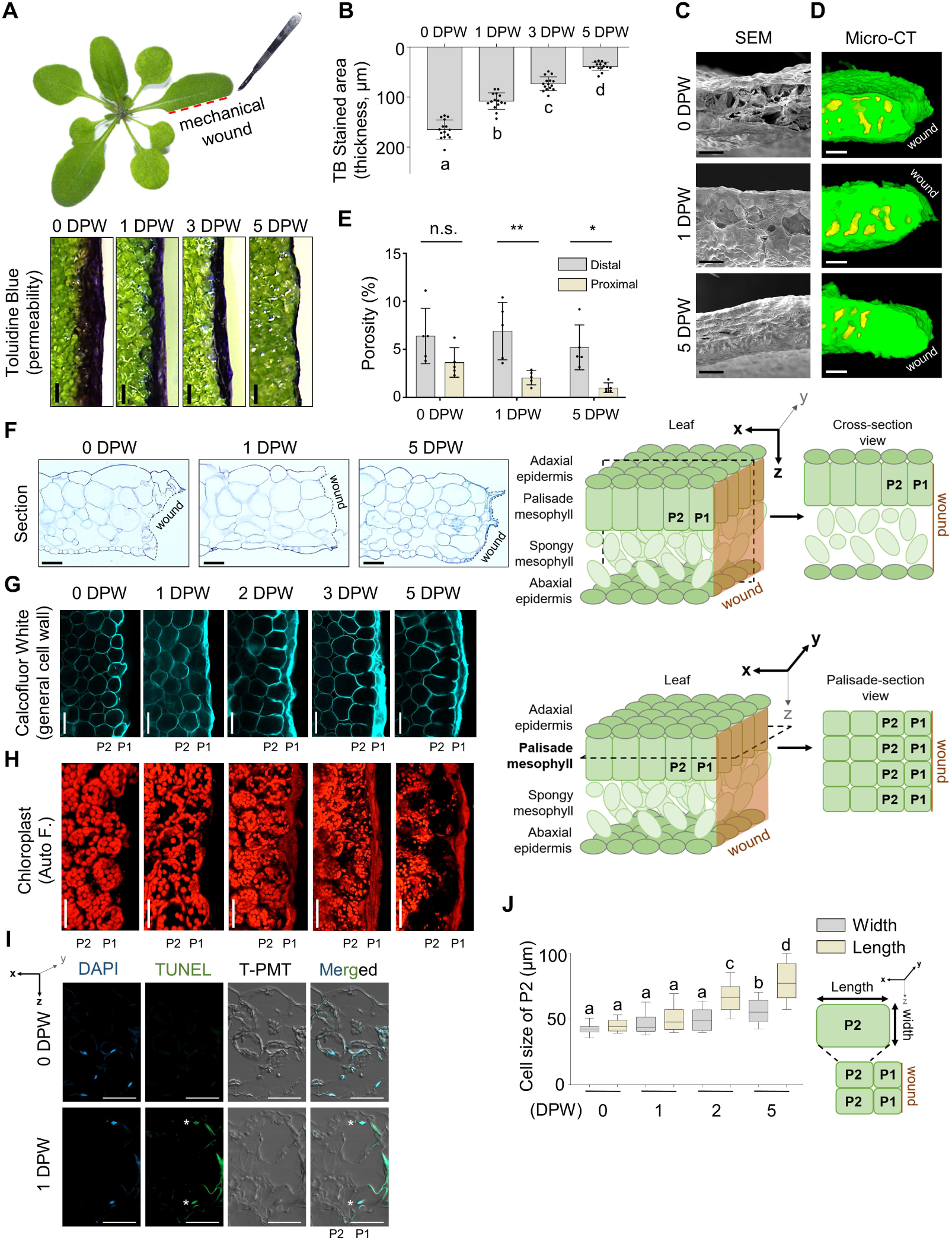
Wounding induces multilayered reorganization of mesophyll cells in mature leaves. (A and B) The permeability of wound sites was assessed using toluidine blue (TB) staining (A), and the stained area was quantified (B). The schematic representation illustrates the leaf wound created by a knife. The error bars indicate mean ± SD—*n* = 15 from 3 independent experiments. See also Quantification and Statistical Analysis in the STAR Methods. (C) Scanning electron micrographs of the wounded leaf show the gradual sealing process of the surface exposed to injury at 0, 1, and 5 DPW. (D and E) Changes in cellular architecture were measured using micro-CT. Three-dimensional-rendered transverse section images of a wounded leaf obtained via micro-CT show leaf tissues in green and airspaces in yellow (D). Porosity was quantified for *n* = 5 samples (E). Proximal and distal porosity was quantified at distances of 0–150 μm and 150–300 μm from the wounded site, respectively. The error bars indicate mean ± SD. See also Figure S1. (F) Sectioned images of the wounded leaf at 0, 1, and 5 DPW. The schematic image illustrates the orientation of the obtained sections. (G) Confocal micrographs display the morphological changes in palisade mesophyll cells in response to wounding. Calcofluor-white was utilized to visualize the cell walls. A schematic image illustrates the orientation of the optical view used for the confocal micrographs. (H) Maximum projection images show the chloroplast autofluorescence of P1 and P2. (I) Confocal micrographs display TUNEL staining of wounded leave sections. White symbol indicates TUNEL-positive P1 cells at 1 DPW. (J) Quantification of P2 cell length. A schematic image illustrates the directions for the width and length. The error bars indicate mean ± SD—*n* = 100. In (B) and (J), statistical significance was analyzed using one-way ANOVA, followed by Tukey’s post-hoc test. Data points with different letters indicate significant differences represented by p < 0.05. In (E), statistical significance was determined by Student’s t-test (*p < 0.05, **p < 0.01, n.s. = no significance). Auto F, autofluorescence; P1, protective layer 1; P2, protective layer 2. Scale bars, 100 μm (A, C), 50 μm (D, F, G, H).

To investigate further the morphological changes in palisade mesophyll cells during wound healing, we stained the cell walls with the fluorochrome Calcofluor-white and observed the tissue by confocal microscopy (Figure 1G). Immediately after the injury, we saw cell debris from the ruptured cells at the outer edge of the wound, and intact layers of cells beneath. The first intact mesophyll cell layer exposed by the wound we designated protective layer 1 (P1), and the second protective layer 2 (P2). At 1 DPW, cell wall material had accumulated over P1, and this cell layer had begun to shrink significantly. By 3 DPW, there was substantial loss of cellular structure and loss of autofluorescence from chloroplasts in P1 (Figure 1G and 1H). Terminal deoxynucleotidyl transferase dUTP nick-end labeling (TUNEL) analysis (Gorczyca et al., 1993; Tripathi et al., 2017) revealed that this cellular disintegration in P1 was accompanied by DNA breakage, indicating a process of programmed cell death (PCD) (Figure 1I). Concurrent with cell death in P1, cells in P2 elongated transversely toward the wound site, eventually organizing into aligned rows (Figure 1G and 1J). Together, these data point to spatial and temporal coordination between P1 and P2 during the wound healing process.

### Changes to the transcriptome during wound healing

To better understand how wounded mesophyll cells rearrange and subsequently form a surface barrier, we analyzed transcriptome profiles at 12 time points over five days following mechanical wounding in 4-week-old *Arabidopsis* leaves. Principal component analysis demonstrated the change in transcriptome profiles over time (Figure S2A). To identify genes whose expression correlated at various time points following wounding, we used weighted gene co-expression network analysis (Langfelder and Horvath, 2008) and identified 23 modules of co-expressed genes (Figures 2A and S2; Table S1). Modules with the most co-expressed genes were predominantly enriched with genes expressed at specific time points after wounding, suggesting that transcriptome profiles reflect temporal changes post-wounding (Figure 2A). To elucidate the characteristics of each module, we analyzed gene ontology terms. Since wound healing involves reorganization of cells and remodeling of their cell walls, we concentrated on five modules enriched for genes related to hormone signaling and cell wall dynamics, which we designated wound healing modules (WMs) 1–5 (Figure 2B; Table S1). WM1, which shows increased expression 1–2 hours post-wounding (HPW), was enriched notably in genes related to responses to hormones, including JA and ethylene (Figure 2B). Conversely, WM2, WM3, WM4 and WM5 were expressed at later time points, and were enriched predominantly in genes involved in cell wall remodeling, including those related to the biosynthesis of surface lipids and lignin (Figure 2B). Further analysis of genes associated with phytohormones and cell wall remodeling revealed that most were enriched at the time points represented by the modules, although expression of a small subset of these genes was more variable (Figure 2C–E). Ethylene-related genes were predominantly expressed before 12 HPW, and subsequently upregulated after 1 DPW. Surface lipid synthesis genes exhibited distinct expression patterns in early (before 1 DPW), intermediate (between 1 and 2 DPW), and late (after 3 DPW) phases, and lignin-related genes were expressed in the early and late phases. These data highlight the multifaceted nature of cell wall remodeling during wound healing, which is temporally regulated with distinct phases characterized by expression of specific genes.

**Figure 2.**
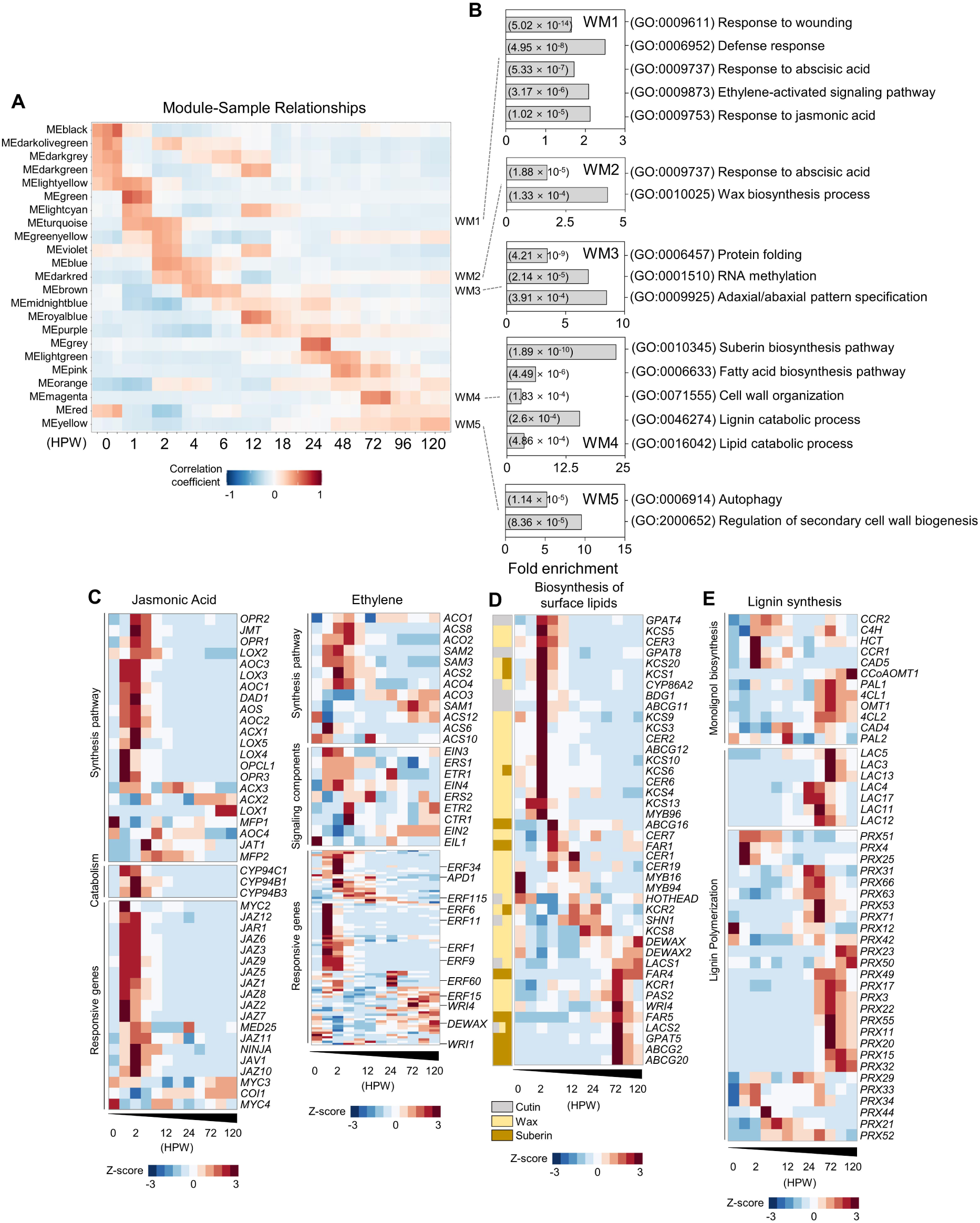
Changes to the transcriptome during wound healing. (A) To identify highly correlated genes at different time points following wounding, weighted gene co-expression network analysis (WGCNA) was performed using a soft-thresholding power of β□=□6. See also Figure S2. (B) Gene Ontology analysis of wound healing modules (WMs). The box indicates the fold enrichment of GO and the value in the box indicates the P-value of each GO terms. See also Table S1. (C–E) Heatmap showing expression patterns of the genes related to the jasmonic acid and ethylene pathway (C), biosynthesis of surface lipids (D), and lignin biosynthesis (E). See also Table S2.

### ATML1 mediates differentiation of epidermal cells to seal the wound surface

We observed suberin, a cell wall component involved in wound healing (Serra et al., 2022), deposited at wound sites (Figure S3A). Our RNA–seq data revealed that the genes related to suberin synthesis and those related to cuticular wax synthesis were both induced by wounding (Figure 2D), however, the genes that synthesize cutin, the waxy polymer that is the main component of the cuticle, were induced much sooner than suberin was deposited (Figure S3A and S3B). We employed transmission electron microscopy to examine deposition of the protective layer and changes to the cell wall at the wound site at high resolution, and observed accumulation of electron-dense substances on the outer surfaces of the cell walls (Figure 3A). Initially appearing as small spots, these substances gradually elongated and eventually covered the entire surface, providing a continuous protective layer (Figure 3A).

**Figure 3.**
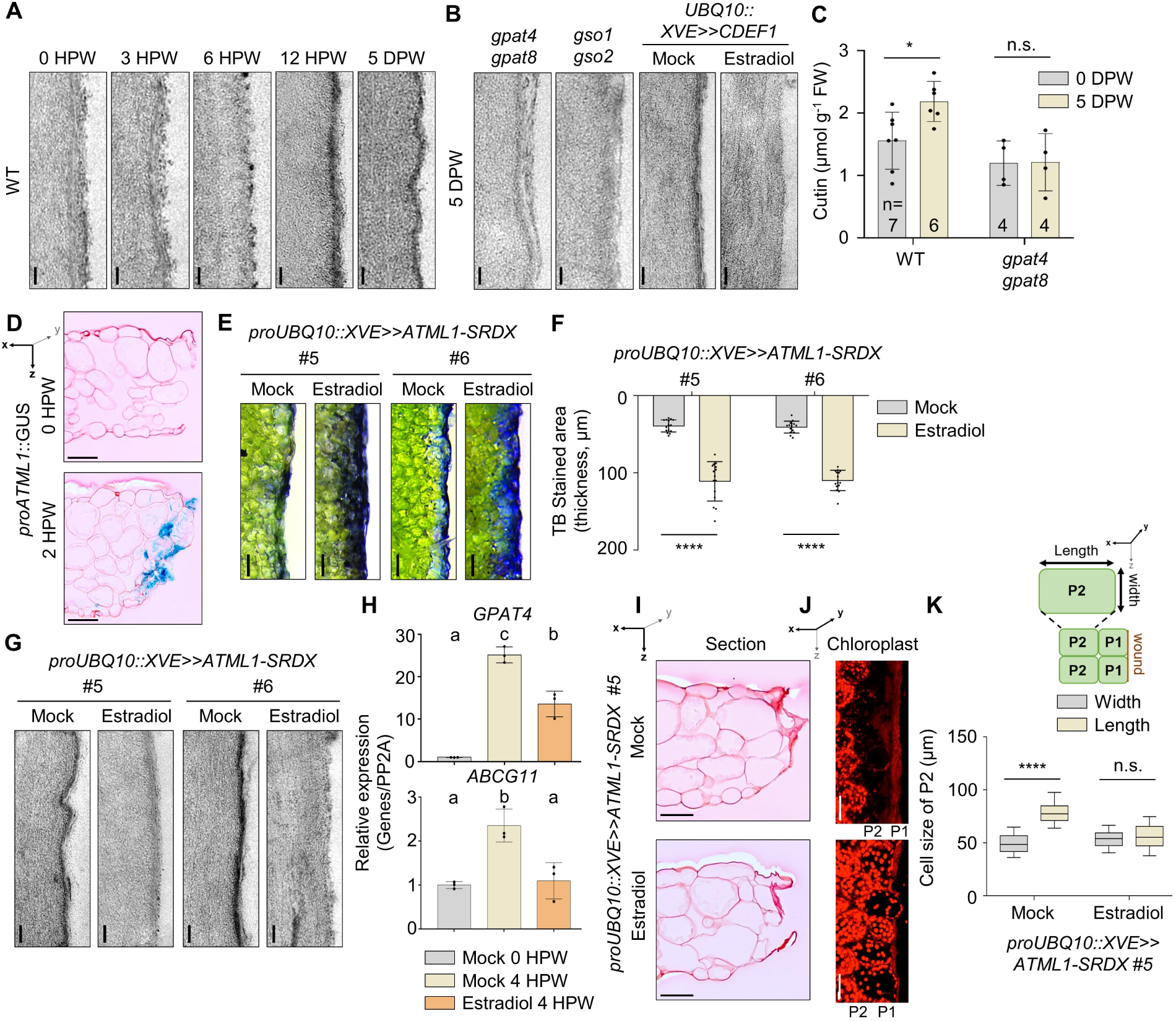
The wound surface is sealed by an ATML1-mediated cuticle at P1. (A and B) Transmission electron micrographs of P1 showing the accumulation pattern of electron-dense material outside the cell wall in the WT (A), various cuticle deficient mutants, and transgenic plants expressing *proUBQ10::XVE-CDEF1* (B). See also Figure S3C. (C) Quantification of total cutin composition in wounded leaves of WT and *gpat4 gpat8* mutants. The error bars indicate mean ± SD. See also Figure S3D. (D) Sectioned images of wounded leaves show the expression of *ATML1* in P1 in transgenic plants harboring *proATML1::GUS* at 2 HPW. GUS stained cell was visualized with blue clour, and Sapranin staining was used to visualize cell morphology. See also Figure S3E. (E and F) Toluidine blue (TB) staining shows the permeability of transgenic plants expressing *proUBQ10::XVE>>ATML1-SRDX* in the absence (mock) or presence of 1 μM estradiol (E), along with quantification of TB-stained area (F). The error bars indicate mean ± SD—*n* = 15 from 3 independent experiments. (G) Transmission electron micrographs of cuticle structure at 5 DPW in transgenic plants expressing *proUBQ10::XVE>>ATML1-SRDX* in the absence (mock) or presence of 1 μM estradiol. (H) RT-qPCR analysis showing expression of *GPAT4* and *ABCG11* in *proUBQ10::XVE>>ATML1-SRDX* in the absence (mock) or presence of 1 μM estradiol at 4 HPW. Expression values normalized to *PP2A* gene levels were displayed as relative to 0-hour mock control—*N* = 3 biological replicates. (I–K) Wound-induced cellular reorganization at 5 DPW was shown with sectioned images with Safranin staining (I), maximum projection images of confocal microscopy displaying chloroplast autofluorescence (J), and expansion of P2 cells (K). Images and data were obtained from *proUBQ10::XVE>>ATML1-SRDX* #5 in the absence (mock) or presence of 1 μM estradiol. The error bars indicate mean ± SD—*n* = 100 cells. In (C), (F), and (K), statistical significance was determined by a Student’s t-test (*p < 0.05, ****p < 0.0001). In (H), statistical significance was determined by a one-sided Kruskal–Wallis test with Holm correction, followed by Dunn’s post-hoc test. Data points with different letters indicate significant differences representative of p < 0.05. Scale bars, 100 nm (A, B, G), 100 μm (E), 50 μm (D, I, J).

Suberin typically accumulates between the cell wall and the plasma membrane (De Bellis et al., 2022; Serra and Geldner, 2022), whereas we found the electron-dense substances accumulated outside the cell wall (Figure 3A), where one would expect to find cuticle (Arya et al., 2021). To investigate whether the electron-dense substance might be cuticle, we analyzed the mutant plants *gpat4 gpat8* and *gso1 gso2*, which have abnormal cuticle formation (Li et al., 2007; Tsuwamoto et al., 2008). The leaves of these plants did not accumulate the electron-dense substance after wounding (Figure 3B). Moreover, transgenic plants overexpressing *CUTICLE DESTRUCTION FACTOR 1* (*CDEF1*), a gene that encodes a lipolytic enzyme of the Gly-Asp-Ser-Leu (GDSL) lipase/esterase protein family that degrades cuticle (Takahashi et al., 2010), accumulated much less of the electron-dense substance after wounding (Figure 3B). This correlated with increased permeability of the leaves in the mutant plants, as revealed by staining with toluidine blue (Figure S3C). Gas chromatography-flame ionization detection (GC–FID) analysis revealed a significant increase in total cutin content during the wound healing process, which was not observed in the *gpat4 gpat8* mutants (Figures 3C and S3D). The predominant cuticle monomer in the healed area was α,ω-octadecadiendioic acid (C18:2 DCA) (Figure S3D), a fatty acid component of leaf cuticle (Berhin et al., 2019; Coen et al., 2019). Collectively, our results demonstrate that the wound-healing process in mature *Arabidopsis* leaves involves cuticle synthesis. This challenges the traditional view of the cuticle simply as an epidermal barrier that, unlike suberin, is not involved in wound healing.

Cuticle accumulation is a well-established marker of epidermis formation (Delude et al., 2016; Kunst and Samuels, 2003; San-Bento et al., 2014). Thus, our findings suggested that wound-healing process in mature leaves might involve differentiation of exposed mesophyll cells into epidermal cells. Indeed, two transcription factors that regulate the differentiation of epidermal cells, *ARABIDOPSIS THALIANA MERISTEM L1* (*ATML1*) and *PROTODERMAL FACTOR2* (*PDF2*), were induced in P1 cells at 2 HPW (Figures 3D and S3E). ATML1 and PDF2 play redundant but critical roles in epidermal specification during embryogenesis (Abe et al., 2003; Ogawa et al., 2015; Takada et al., 2013). To investigate whether wound healing involves differentiation of mesophyll cells into epidermal cells, we expressed a gene encoding ATML1 fused to the transcriptional repressor domain SRDX, under the control of estradiol (Iida et al., 2019) (Figure S3F and S3G). The ATML1–SRDX fusion protein acts as a dominant repressor of ATML1, thus inhibiting differentiation of epithelial cells. Treatment of *ATML1–SRDX* mutant seedlings with estradiol resulted in pronounced abnormalities in epidermal differentiation (Figure S3H). Wounds in the adult leaves of *ATML1–SRDX* mutants, remained permeable 5 DPW (Figure 3E and 3F), and the formation of a protective cuticle on the injured surface was severely impaired (Figure 3G). Wounded *ATML1–SRDX* mutant plants induced with estradiol did not increase expression of genes associated with cutin biosynthesis or transport, including *GPAT4* and *ABCG11* (Li et al., 2007; McFarlane et al., 2010), as much as the control plants (Figure 3H). Also, the reorganization of cell layers and chloroplast degradation seen in the controls were inhibited in the presence of estradiol (Figure 3I–K). These findings emphasize the critical role of ATML1 in healing wounds in mature leaves by converting mesophyll cells in P1 into epidermal cells and triggering the deposition of a protective layer of cuticle.

### Ethylene–RbohE signaling is crucial for P1 cell death and cuticular wax formation

After ATML1-mediated cuticle formation by newly differentiated epidermal cells, P1 undergoes programmed cell death and P2 takes its place just beneath the cuticle (Figure 1G and 1I). To understand the molecular mechanisms of cell death in P1, we examined our transcriptomics data for expression of genes associated with programmed cell death. Reactive oxygen species (ROS) serve as secondary messengers, regulating diverse developmental and signaling processes, including cell death (Gechev et al., 2006; Mittler et al., 2022). NADPH oxidases, known as respiratory burst oxidase homologs (RBOHs) in plants, are the key enzymes that generate ROS (Sagi and Fluhr, 2006). Intriguingly, the RBOH genes *RbohC, RbohD, and RbohF* were expressed in the early phase, whereas *RbohE* and *RbohF* were expressed in the late phase of wound healing (Figure 4A). This two-phase pattern of expression was reflected in the accumulation of ROS at the wound sites in wild-type and *rbohD* and *rbohE* plants 1 HPW and 3 DPW (Figure S4A), indicating that *RbohD* is involved at early stages and *RbohE* at later stages of wound healing.

**Figure 4.**
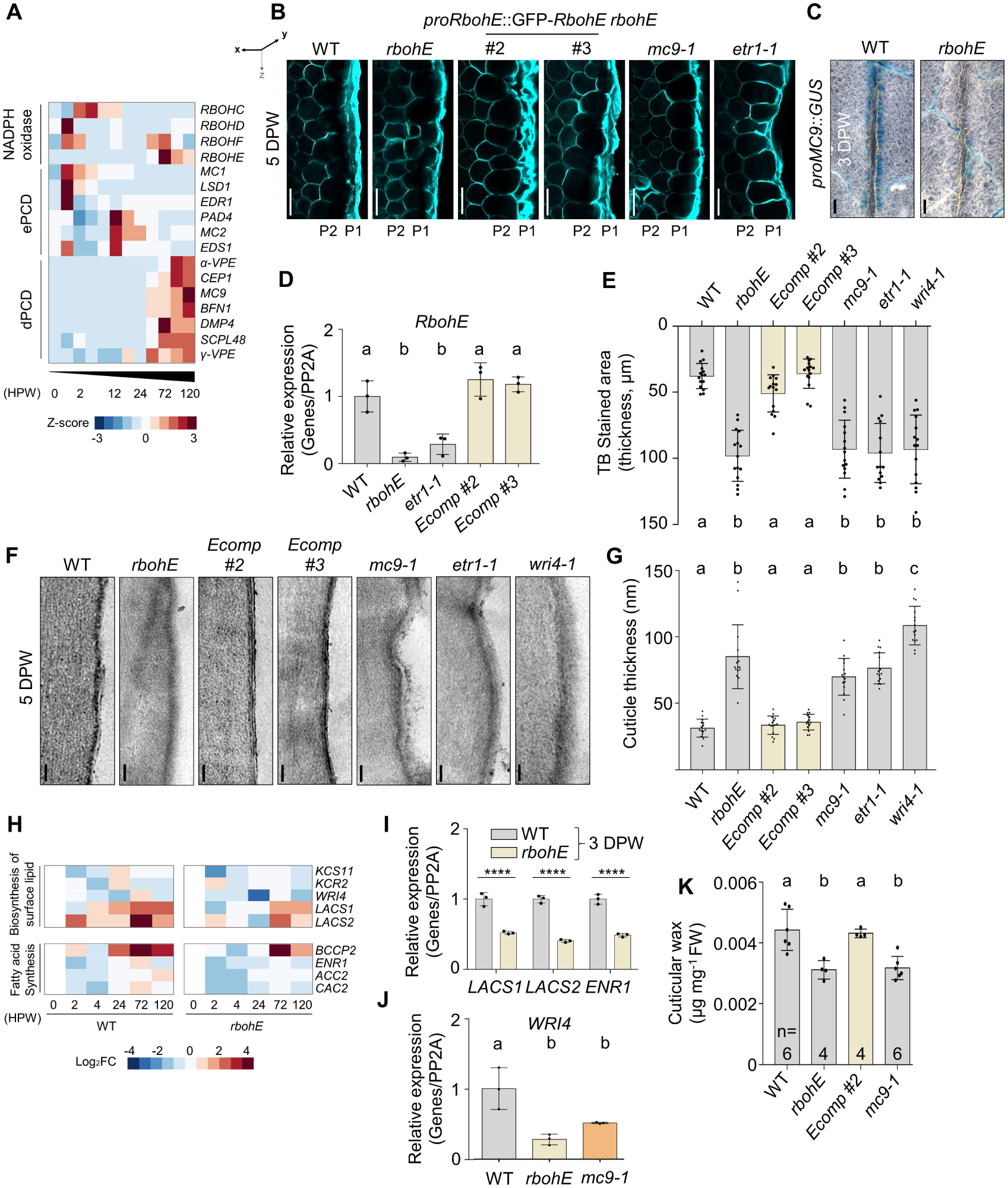
Ethylene–RbohE signaling is crucial for P1 cell death and cuticular wax formation. (A) Heatmap showing expression patterns of genes encoding NADPH oxidases and genes associated with environmental programmed cell death (ePCD) or developmental PCD (dPCD). See also Table S2. (B) Confocal micrographs showing the difference in P1 cell death in WT and various mutants after calcofluor-white was used to visualize the cell wall. See also Figure S4B and S4C. (C) Promoter-GUS analyses of *MC9* in WT and *rbohE* at 3 DPW. (D) Wound-induced *RbohE* expression in WT and *rbohE, etr1-1*, and *rbohE* complemented with *proRbohE::GFP-RbohE* (Ecomp) at 3 DPW—*N* = 3 biological replicates. (E) Quantification of toluidine blue (TB) staining in WT and various mutants. The error bars indicate mean ± SD—*n* = 15 from 3 independent experiments. (F and G) Transmission electron micrographs of the cell wall at P1 in various mutants (F), along with quantification of the thickness of the cuticular wax layer (G). The error bars indicate mean ± SD—*n* = 15 from 3 independent experiments. See also Quantification and Statistical Analysis in the STAR Methods. (H) The heat map shows expression patterns of wax synthesis genes and fatty acid synthesis genes in WT and *rbohE*, and it is presented with a Log_2_FC scale relative to the 0-hour control for each background. Downregulated genes in *rbohE* (Log_2_FC difference > 0.5 at 1 or 3 DPW) were represented in the heat map. See also Figure S4D, S4E and Table S3. (I) Expression of the wax-related genes *LACS1, LACS2* and *ENR1* was analyzed by RT-qPCR in WT, *rbohE* at 3 DPW. The expression values were normalized to the PP2A gene, and the fold change was calculated relative to the 0-hour control for each background. The expression levels are presented relative to the WT. The error bars indicate mean ± SD—*N* = 3 biological replicates. (J) *WRI4* expression in WT*, rbohE* and *mc9-1* at 3 DPW. The expression values were normalized to the PP2A gene, and the fold change was calculated relative to the 0-hour control for each background. The expression levels are presented relative to the WT. The error bars indicate mean ± SD—*N* = 3 biological replicates. (K) Quantification of total cuticular wax composition in wounded leaves at 5 DPW. The error bars indicate mean ± SD. See also Figure S4G. In (D), (J) and (K), statistical significance was determined by a one-sided Kruskal–Wallis test with Holm correction, followed by Dunn’s post-hoc test. In (E) and (G), statistical significance was analyzed using one-way ANOVA, followed by Tukey’s post-hoc test. In (I), statistical significance was determined by a Student’s t-test (***p < 0.001, ****p < 0.0001). Data points with different letters indicate significant differences representative of p < 0.05. Scale bars, 50 μm (B), 100 μm (C), 100nm (F).

Interestingly, cell death in the P1 layer was suppressed in *rbohE* mutants, and this effect was reversed when the *RbohE* gene was complemented (Figures 4B, S4B), indicating a crucial role for RbohE in cell death in this layer. *RbohE* expression coincided with expression of genes related to developmentally regulated PCD (Figure 4A). Mutation of one of these developmental PCD genes, *MC9*, also suppressed PCD in P1 (Figure 4B). By contrast, mutation of three genes related to environmentally induced PCD, *MC1*, *MC2,* and *LSD1*, had little or no effect (Coll et al., 2010) (Figure S4C). Expression of *MC9* at the wound site was significantly suppressed in *rbohE* mutants (Figure 4C), indicating that RbohE-mediated signaling regulates expression of genes involved in cell death.

Ethylene acts upstream of developmental PCD (Huysmans et al., 2017; Rantong et al., 2015; Völz et al., 2013). Thus, we hypothesized that ethylene plays a role in controlling PCD upstream of RbohE during wound healing in mature leaves. Indeed, cell death in P1 was suppressed in the ethylene signaling mutant, *etr1-1* (Figure 4B), and *RbohE* was downregulated in this mutant (Figure 4D). When cell death in P1 was suppressed, wound healing was impaired, as indicated by permeability to toluidine blue (Figure 4E) and transmission electron microscopy at 5 DPW (Figure 4F and 4G). These findings suggest that ethylene and RbohE regulation of developmental PCD genes is crucial for death of the cells in P1.

Not only did the wound surface remain permeable in the *rbohE* mutant, but also formation of the cuticle layer was compromised at 5 DPW, with a blurred and less compact cuticle than that seen in the wild type (Figure 4F and 4G). Expression of genes involved in cutin synthesis increased predominantly within 2–6 HPW – preceding expression of *RbohE* – but no significant differences were observed in expression of these genes in the *rbohE* mutants when compared with wild-type plants (Figure S4D and S4E), and a cuticle barrier formed at 12 HPW in *rbohE* mutants like that in the wild type (Figure S4F). These results suggest that additional processes, potentially mediated by RbohE, are required for the maturation of the cuticle layer.

The cuticle layer, composed of cutin and cuticular wax, relies on a delicate balance between these components, with cuticular wax playing a crucial role in ensuring proper formation of the cuticle (Qin et al., 2011; Suh et al., 2005). Given that many genes associated with wax biosynthesis are expressed at the same time as *RbohE* at 2–3 DPW, we hypothesize that RbohE mediates cuticular wax synthesis. Indeed, we observed that genes involved in fatty acid elongation for wax biosynthesis, such as *KCR2, WRI4*, *LACS1,* and *LACS2* (Beaudoin et al., 2009; Kim et al., 2017; Lu et al., 2009; Park et al., 2016; Schnurr et al., 2004) were downregulated in *rbohE* mutants (Figure 4H-J), and defects in wax formation post-wounding were also observed (Figures 4K and S4G). Genes downregulated by *rbohE* included not only those involved in fatty acid elongation but also those involved in fatty acid biosynthesis such as *BCCP2*, *ENR1*, *ACC2* and *CAC2* (Figure 4H and 4I). Interestingly, most of these genes are regulated by a transcriptional activator WRI4 (Park et al., 2016). In a *WRI4*-deficient mutant, we observed significant defects in cuticular layer formation and surface sealing after wounding (Figure 4E–G), mirroring the *rbohE* mutant phenotype. This suggests that WRI4 plays a key role in RbohE-mediated wax formation. Furthermore, we also observed downregulation of *WRI4* and defects in wax formation post-wounding in *mc9-1* mutants (Figures 4J, 4K and S4G). Considered together, our evidence that RbohE is required both for expression of developmental PCD genes and cuticular wax biosynthesis suggests that the sacrifice of P1 cells is crucial in providing sufficient wax to the surface, ultimately leading to the completion of the protective layer.

### JA–RbohD signaling regulates the lignification of P2

The permeability of the wound to toluidine blue indicates that the barrier continues to be reinforced from 3–5 DPW following P1 cell death, likely involving additional cell wall remodeling in P2 (Figure 1B). To understand the role of P2 in wound healing, distinct from cuticle formation, we focused on lignin deposition – another cell wall remodeling process. We used basic fuchsin to stain the lignin in cleared tissue (Ursache et al., 2018) of wounded leaves from wild-type plants and observed clear lignin deposition around P2 cells at 5 DPW (Figure S5A). Lignin biosynthesis comprises two stages: monolignol biosynthesis and polymerization of secreted monolignols (Dixon and Barros, 2019). Our transcriptomics data revealed a notable time gap between these stages: expression of monolignol biosynthesis genes was significantly enhanced before 12 HPW, whereas expression of genes related to polymerization of monolignols, such as *Peroxidases* and *Laccases,* was enhanced from 2–5 DPW (Figures 2E and S5B). This two-phase expression pattern suggests that, although lignin formation is observed at 5 DPW, signaling for precursor production must begin early on.

The phytohormone JA accumulates rapidly in tissues proximal and distal to plant injury sites and stimulates the wound response (Koo and Howe, 2009; Mousavi et al., 2013). In *aos* and *myc2 myc3 myc4* mutants, in which the JA synthesis and signaling pathways are blocked, respectively (Fernandez-Calvo et al., 2011; Park et al., 2002), permeability of the wound site to toluidine blue was greater than in wild-type plants (Figure 5A), suggesting a role for JA in local wound healing; however, in the *aos* mutant, which is defective in JA synthesis, cuticle deposition in P1 was remarkably similar to that in the wild type (Figure 5B). By contrast, lignin deposition, as indicated by basic fuchsin staining, was markedly diminished in the *aos* and *myc2 myc3 myc4* mutants, and treatment of the *aos* mutant with methyl jasmonate, a volatile form of JA, restored it to WT levels (Figure 5C and 5D). Interestingly, lignin deposition was also restored in the *aos* and *myc2 myc3 myc4* mutants by treatment with monolignol (Figure 5C and 5D), indicating that these mutants remain able to polymerize lignin when the monomer is supplied. This suggests that JA may regulate lignin formation by modulating the expression of monolignol synthesis genes, a hypothesis confirmed by RT–qPCR analysis, showing downregulation of these genes in the *aos* and *myc2 myc3 myc4* mutants when compared with the wild type (Figures 5E and S5C).

**Figure 5.**
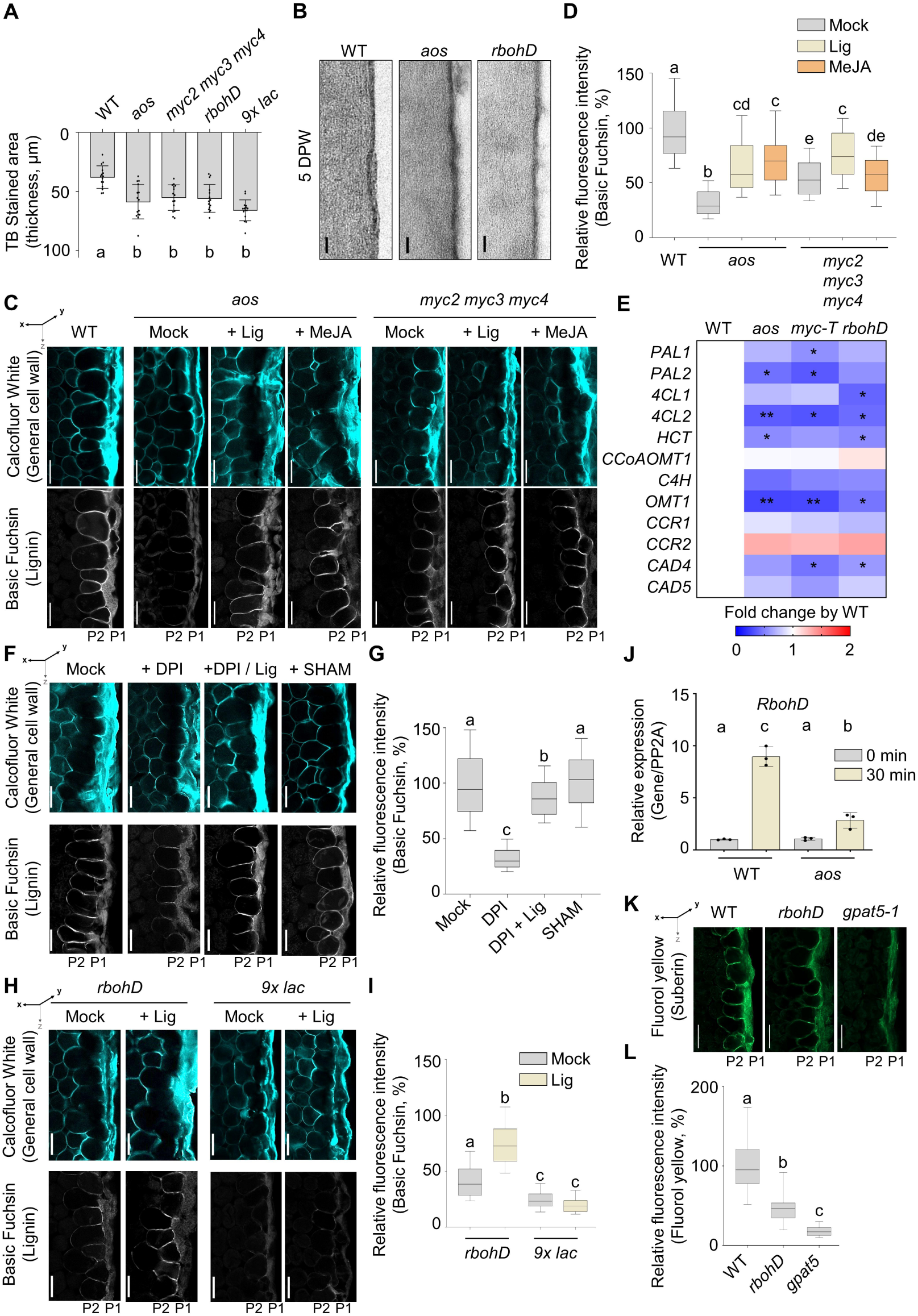
JA–RbohD signaling regulates the lignification of the P2 layer. (A) Quantification of toluidine blue (TB) staining in WT and various mutants. The error bars indicate mean ± SD—*n* = 15 from 3 independent experiments. (B) Transmission electron micrographs of cuticle layer formation in WT, aos and *rbohD* at 5 DPW. (C and D) Confocal micrographs show the P2 lignification in WT and JA-related mutants at 5 DPW (C). A total of 30 µM coniferyl alcohol and 30 µM sinapyl alcohol (Lig) or 10 µM Methyl Jasmonate (MeJA) were infiltrated before mechanical wounding. Calcofluor-white was used to visualize the cell wall, while basic fuchsin was employed for lignin detection. Lignin accumulation was quantified using the fluorescence signal of basic fuchsin, with values expressed relative to the WT (D). *n* = 100. See also Quantification and Statistical Analysis in the STAR Methods. (E) A heatmap showing the expression of monolignol biosynthesis-related genes in WT and various mutants at 4 HPW. Expression levels were assessed using RT-qPCR, normalized to PP2A expression, and presented relative to WT. The mean values of 3 biological replicates were represented in heat map. *myc-T, myc2 myc3 myc4*. See also Figure S5C. (F–G) Confocal micrographs show the lignification at P2 after basic fuchsin staining (F). A total of 100 μM diphenyleneiodonium (DPI), 50 μM salicylhydroxamic acid (SHAM), or 30 µM coniferyl alcohol and 30 µM sinapyl alcohol (Lig) were infiltrated before mechanical wounding. Lignin accumulation was quantified using basic fuchsin fluorescence intensity relative to the WT mock control (G). *n* = 100. (H-I) Confocal micrographs show the lignification at P2 of *rbohD* and *9x lac*. 30 µM coniferyl alcohol and 30 µM sinapyl alcohol (Lig) or Mock control were infiltrated before mechanical wounding (H). Lignin accumulation was quantified using basic fuchsin fluorescence intensity relative to the WT mock control (I). *n* = 100. (J) Expression of *RbohD* in WT and *aos* at 30 minutes post-wounding was assessed using RT-qPCR. Expression levels were normalized to *PP2A* and presented relative to the 0-hour control for each background. The error bars indicate mean ± SD—*N* = 3 biological replicates. See also Figure S5D. (K and L) P2 suberization was visualized using Fluorol Yellow staining and confocal microscopy in WT, *rbohD*, and *gpat5* mutants at 5 DPW (K). The fluorescence intensity was quantified and presented relative to WT (L). *n* = 100. See also Quantification and Statistical Analysis in the STAR Methods. In (A), (D), (G), (I), and (L), statistical significance was analyzed using one-way ANOVA, followed by Tukey’s post-hoc test. In (J), statistical significance was determined using a one-sided Kruskal– Wallis test with Holm correction, followed by Dunn’s post-hoc test. Data points with different letters indicate significant differences representative of p < 0.05. The box and whisker plot (D, G, I, L) shows data between the 10th and 90th percentiles. The box represents the interquartile range, with the line inside the box indicating the median. Scale bars, 100 nm (B), 50 μm (C, F, H, K).

Previous research indicates that ROS generated by RBOHs contribute to lignin biosynthesis by acting as signaling mediators and/or by providing substrates necessary for peroxidase-mediated lignin polymerization (Barros et al., 2015; Lee et al., 2013; Lee et al., 2018). Treatment with the RBOH inhibitor diphenyleneiodonium chloride inhibited lignin formation in response to wounding, whereas co-treatment with the inhibitor and monolignols restored lignin production (Figure 5F and 5G), suggesting a role for RBOHs in monolignol biosynthesis during wound healing. *RBOHD* was expressed at 1–2 HPW, slightly before the monolignol biosynthesis-related genes (Figures 2E and 4A), and lignin deposition was significantly reduced in the *rbohD* mutant (Figure 5H and 5I). Since the peroxidase inhibitor salicylhydroxamic acid did not affect lignification (Figure 5F and 5G), and monolignol treatment increased lignification in the *rbohD* mutant (Figure 5H and 5I), we postulated that the primary role of RbohD-mediated ROS accumulation, like that of JA, is in monolignol biosynthesis. Wound-induced expression of monolignol biosynthesis genes was impaired in the *rbohD* mutant (Figures 5E and S5C). Moreover, the nonuple laccase mutant *9x lac* (mutant in *lac1, lac3, lac5, lac7, lac8, lac9, lac12, lac13 and lac 16*), which is defective in the enzymes that promote polymerization of monolignols into lignin, lacked wound-induced lignin and did not respond to monolignol treatment (Figure 5H and 5I). These data suggest that JA and RbohD may function in the same pathway. Indeed, *RbohD* expression was significantly reduced (Figure 5J), and ROS accumulation was diminished in the *aos* mutant (Figure S5D), indicating that JA signaling promotes monolignol synthesis through RbohD-mediated ROS signaling. Suberin deposition was also significantly impaired in the *rbohD* mutant (Figure 5K and 5L), whereas cuticle deposition in P1 proceeded normally (Figure 5B). Together with the increased permeability in the *rbohD* mutant (Figure 5A), these findings highlight the importance of ligno-suberization in sealing the wound surface below the cuticular wax barrier.

### ATML1 regulates the specification and function of cells in P2

Our observations reveal that whereas P1 serves initially as the primary protective layer by forming a cuticle, the later stage of fortification, marked by ligno-suberization, is driven by P2, which assumes the frontline role following the death of the cells in P1. This raises the question of whether the cells in P2, like those in P1, differentiate into epidermal cells and, if so, whether ATML1 mediates this process. *ATML1* expression was again upregulated at 1–2 DPW (Figures 6A, 6B, and S6A), supporting the hypothesis that ATML1 is involved in the fate of P2 cells. Moreover, induction of the dominant repressor gene *ATML1-SRDX* with estradiol at 12 HPW, after P1 establishment, reduced expression of laccases genes (Figure 6C), and concurrently inhibited lignin formation (Figure 6D). Moreover, we observed a dramatic decrease in chlorophyll content in mature P2 at 5 DPW, consistent with an epidermal character with few or no chloroplasts (Barton et al., 2016; Pyke, 2009), and this decrease was also inhibited by inducing *ATML1-SRDX* with estradiol (Figure 6D). Together, these findings suggest a critical role for ATML1 in determining the fate of P2 and regulating its function. *PDF2*, which is functionally redundant with *ATML1* (Ogawa et al., 2015), was also upregulated at 1–2 DPW (Figure S6B), around the same time as *ATML1*. No significant effects on lignin deposition were observed in the single mutants of either *atml1-3* or *pdf2-4* (Figure S6C and S6D), indicating that the functions of ATML1 and PDF2 in P2 regulation are redundant.

**Figure 6.**
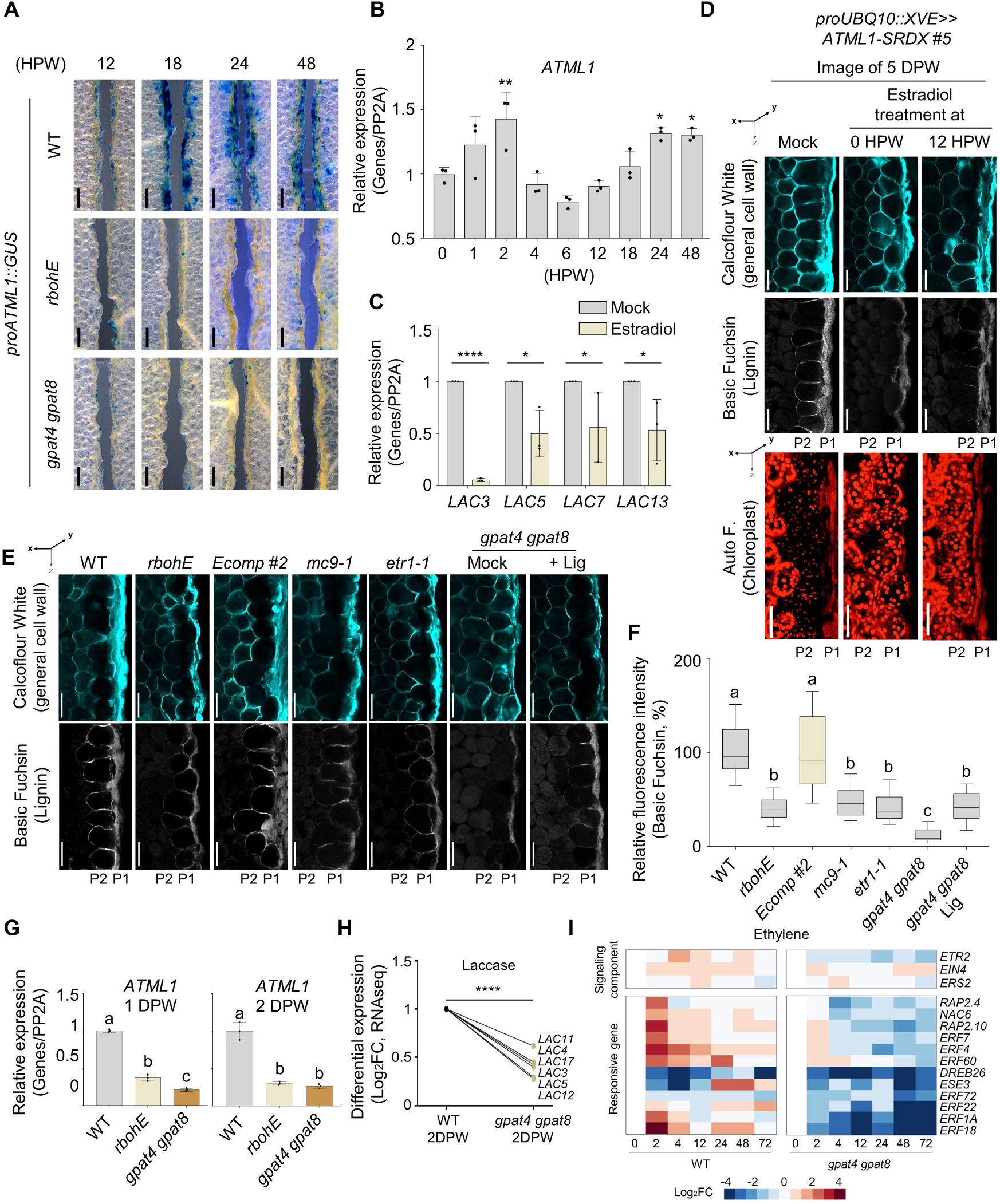
The ATML1 module mediates P2 specification. (A) Promoter-GUS histochemical analyses of *ATML1* in WT, *rbohE, gpat4 gpat8* at 12 HPW to 2 DPW. See also Figure S6A and S6B. (B) Time series *ATML1* expression of WT. *N* = 3 biological replicates for each time point. (C) Laccases expression in *ATML1–SRDX* #5. Estradiol was administered at 12 HPW, and RNA was extracted at 2 DPW. Expression levels were assessed using RT-qPCR, normalized to PP2A expression, and presented relative to each Mock control. The error bars indicate mean ± SD—*N* = 3 biological replicates. (D) Effects of *ATML1–SRDX* induction on P2 specification. Estradiol or Mock treatment was applied at 0 HPW and 12 HPW and analyzed at 5 DPW using confocal microscopy. Calcuflour-white was used to show cell morphology, basic fuchsin for lignin and chloroplast was visulalized by autoflourescence. See also Figure S6C and S6D. (E and F) Confocal micrographs show P2 lignification in WT and various mutants at 5 DPW (E). Lignin intensity was quantified by measuring fluorescence intensity and displayed relative to WT (F). The box and whisker plot shows data between the 10th and 90th percentiles. The box represents the interquartile range, with the line inside the box indicating the median. *n* = 100. See also Figure S6E. (G) *ATML1* expression in WT, *rbohE,* and *gpat4 gpat8* at 1 and 2 DPW. The expression values were normalized to the PP2A gene, and calculated relative to the 0-hour control for each background. The expression levels are presented relative to the WT. The error bars indicate mean ± SD—*N* = 3 biological replicates. (H) The expression patterns of laccase genes at 2 DPW in WT and *gpat4 gpat8* were analyzed using RNAseq. Differential expression level in a graph is presented as Log_2_FC relative to the 0-hour control for each background, and relative to WT. See also Table S4. (I) The expression patterns of ethylene-related genes in WT and *gpat4 gpat8* were analyzed using RNAseq. Expression level in a heatmap is presented as Log_2_FC relative to the 0-hour control for each background. See also Table S4. In (B), (C) and (H), statistical significance was determined by Student’s t-test (*p < 0.05, **p < 0.01, ****p < 0.0001). In (F), statistical significance was analyzed using one-way ANOVA, followed by Tukey’s post-hoc test. In (G), statistical significance was determined by a one-sided Kruskal–Wallis test with Holm correction, followed by Dunn’s post-hoc test. Data points with different letters indicate significant differences representative of p < 0.05. Scale bars, 100 μm (A), 50 μm (D, E).

The death of P1 appears crucial for P2 to assume the role as the first living layer, leading us to posit what would occur if P1 cells remained alive. In *rbohE*, *mc9-1* and *etr1-1* mutants, in which P1 cell death is inhibited, lignin formation (Figure 6E and 6F) and *ATML1* induction at 1 DPW (Figure 6A and 6G) was substantially reduced. In these mutants, we saw some lignification in P1, but not in P2 (Figure 6E and 6F), indicating that lignin polymerization occurs in the living first layer, which maintains its epidermal identity, rather than specifically in P2 cells.

What determines the timing of P1 cell death? Interestingly, the *gpat4 gpat8* double mutant, which is impaired in early cuticle formation, provides some insights into this question. In this mutant, P1 cell death (Figure 6E) and lignin formation (Figure 6E and 6F) were inhibited. Likewise, in plants overexpressing *CDEF1*, the gene that encodes the GDSL lipase enzyme that degrades cuticle, P1 cell death and lignin formation were inhibited (Figure S6E). These phenotypes suggest that cuticle formation is crucial for triggering P1 cell death. Defects in cuticle formation also affected the specification of P2, similarly with other P1 cell death mutants such as *rbohE* or *mc9-1*. Not only was *ATML1* induction reduced at 1 DPW (Figure 6A and 6G), but similarly to the effect observed with estradiol-induced *ATML1-SRDX*, the expression of laccases genes was also reduced in the *gpat4 gpat8* mutant (Figure 6H). Monolignol treatment slightly enhanced lignin synthesis to detectable levels in *gpat4 gpat8* mutant, with lignin pattern observed in P1 (Figure 6E). This suggests that cuticle formation plays a crucial role in triggering P1 cell death, thereby completing the wound-induced protective layer. The reduced expression of ethylene-responsive genes in *gpat4 gpat8* mutants (Figures 6I) suggests that ethylene signaling may mediate the connection between cuticle formation to P1 cell death. Ethylene, generated in response to wounding, may be trapped as the cuticle forms, thereby inducing ethylene responses and determining the precise timing of P1 cell death.

Our findings indicate that wound healing involves integration of the ethylene and JA signaling pathways with ATML1-mediated epidermal cell specification. Together, they coordinate cell layer-specific functions, leading to multilayered cell wall modifications, including cuticular wax formation and ligno-suberization (Figure 7A). This mode of wound healing is not limited to *Arabidopsis* leaves; we observed similar processes in *Nicotiana benthamiana* and *Capsella bursa-pastoris*. In both species, the outer surface was covered with an electron-dense material, likely cuticle, 5 DPW (Figure 7B and 7C) and, although the number of layers differed, the underlying layers were lignified (Figure 7D).

**Figure 7.**
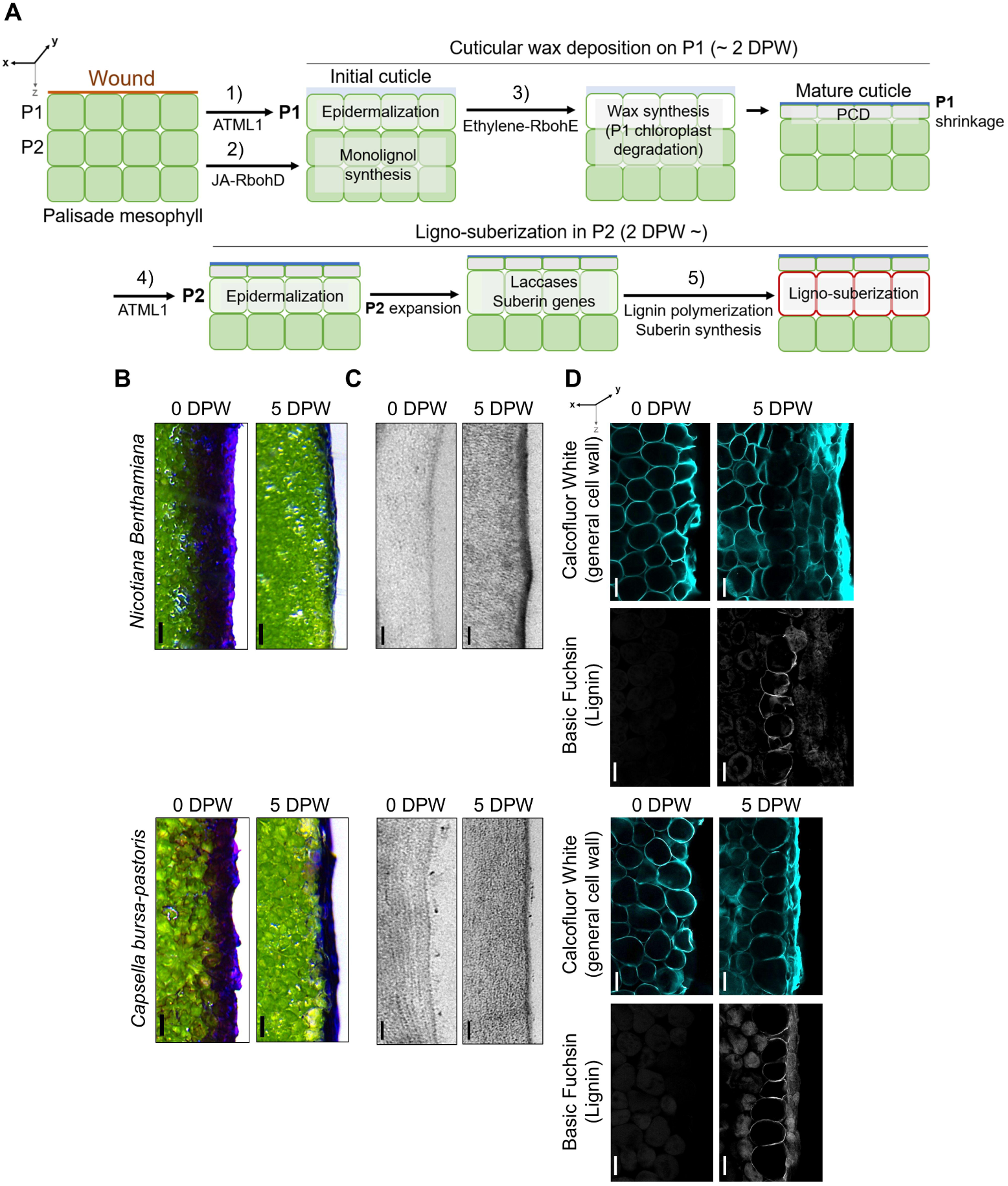
Wound-induced periderm formation in plants. (A) The wound healing process could be divided into the following stages: 1) ATML1-mediated cuticle formation in P1, 2) JA–RbohD-mediated monolignol synthesis, 3) ethylene–RbohE-mediated cell death in P1 and wax synthesis, 4) ATML1-mediated epidermalization of P2, and 5) lignin polymerization and suberization in P2. (B) Image of toluidine blue staining shows the wound barrier formation in *Nicotiana Benthamiana* and *Capsella bursa-pastoris*. (C) Transmission electron micrographs show the wound-induced cuticle layer formation in *Nicotiana benthamiana* and *Capsella bursa-pastoris*. (D) Confocal micrographs show lignification in *Nicotiana benthamiana* and *Capsella bursa-pastoris*. Scale bars, 100 μm (B), 100 nm (C), 50 μm (D).

## DISCUSSION

The epidermis of plant leaves, protected by cuticular wax, is a vital barrier against water loss and various biotic and abiotic threats (Arya et al., 2021; Delude et al., 2016; Kim et al., 2017; Kunst and Samuels, 2003; Suh et al., 2005); however, this barrier is vulnerable to insect attacks and mechanical damage. Here, we investigated the mechanisms that govern wound healing at sites of local physical injury in mature *Arabidopsis* leaves. Our study provides a comprehensive spatiotemporal perspective on the mechanisms that repair the compromised protective layers, notably by forming a multilayered, ligno-suberized barrier coated with cuticular wax – a novel form of wound periderm. This repair process involves two distinct cell layers and is orchestrated by coordination of JA, ethylene and ROS with ATML1-mediated differentiation of mesophyll cells into epidermal-like cells (Figure 7). It differs from wound-periderm formation in tubers, where a suberized barrier forms a seal, and a meristematic cell layer beneath generates suberized cells (Lulai and Neubauer, 2014; Sabba and Lulai, 2002; Schreiber et al., 2005; Serra et al., 2022). This novel epidermalization-based wound healing process occurs in the leaves of diverse plants (Figure 7).

It has been generally accepted that wound healing in plants involves suberin deposition while mesophyll cells maintain their original fate (Sabba and Lulai, 2002; Serra and Geldner, 2022; Serra et al., 2022). Although a recent report demonstrated that the protective layer formed after floral organ abscission in *Arabidopsis* was cuticular (Lee et al., 2018), this remains an exceptional case, and the detailed mechanisms are yet to be elucidated. Our data, which highlight the importance of ATML1-mediated cuticle formation in wound healing, reveal that the epidermalization of mesophyll cells is not an exception confined to floral organ abscission.

Suberin and cuticle are both lipid-based polymers, but they are deposited in different locations: suberin beneath the primary cell wall and cuticle on the outer surface (Arya et al., 2021; De Bellis et al., 2022; Nawrath, 2002). This location of suberin limits its effectiveness as an apoplastic barrier, making the cuticle the more suitable for epidermal protection in organ development and wound healing. Although the detailed mechanisms of epidermal specification are not yet fully understood, it is known that ATML1 and PDF2 act are key transcription factors, directly inducing the expression of cuticle-related genes (Takada, 2013; Takada et al., 2013). The wound-induced expression of *ATML1* and cuticle synthesis-related genes in mesophyll cells suggests that ATML1 also plays a significant role in the epidermalization process during wound healing. So far, elucidating the detail of the epidermalization process has been impeded by lethality issues; however, the wound-healing process presents a promising model to overcome these challenges.

While JA and ROS are known to play a crucial role in the wound response (Denness et al., 2011; Kawano, 2003; Leon et al., 2001; McConn et al., 1997; Miller et al., 2009), their precise functions in local wound healing have remained unclear. Our findings elucidate the mechanisms of action of these factors in space and time during wound healing. The wound-healing process can thus be divided into five stages: ATML1-mediated cuticle formation by P1; JA–RbohD-mediated monolignol synthesis; ethylene–RbohE-mediated cell death and wax synthesis in P1; ATML1-mediated epidermalization of P2, and, finally, lignin polymerization and suberization in P2. Intriguingly, formation of the protective layer occurs in a modular fashion. Cuticular wax formation continues without JA signaling, and lignin formation persists without ethylene signaling. This modular approach to wound healing may reflect an adaptive strategy for managing environmental stresses and pathogen interactions, allowing for cell wall remodeling to adjust to changing conditions and providing alternative pathways if specific signaling routes are blocked by pathogen effectors. Subsequent research would be needed to investigate how pathogens disrupt this process, and to identify the strategies plants use to counteract these disruptions.

We describe how the P1 and P2 layers are coordinated in the wound healing process. The role of P2 becomes relevant only after P1 has died, meaning it is always the outermost living layer that actively forms the protective barrier. When previously internal mesophyll cells are exposed to the outside upon injury, various mechanisms may help them sense this exposure. Notably, changes in mechanical pressure caused by the rupture of surrounding cells might be a significant factor, as demonstrated in wound healing in roots (Hoermayer et al., 2024; Marhava et al., 2019). Recent findings indicate that mechanical pressure reduces *ATML1* expression in mesophyll cells following removal of the epidermis from *Arabidopsis* leaves (Iida et al., 2023). This suggests a relationship between mechanical pressure and *ATML1* regulation. Mechanical pressure alone may not be sufficient, however, to distinguish between changes in mechanical force due to internal cell death and those associated with external exposure due to injury. Changes in the concentrations of substances such as oxygen, volatile chemicals, and humidity, which differ in the internal and external environments of normal leaves, might indicate exposure to the external environment.

Although ATML1 was identified as a master regulator of epidermal layer specification over a decade ago (Abe et al., 2003; Iida et al., 2019; Lu et al., 1996; Ogawa et al., 2015; Peterson et al., 2013; San-Bento et al., 2014; Sessions et al., 1999; Takada et al., 2013), the regulatory mechanisms controlling its expression remain largely unexplored. Our findings reveal two distinct contexts for *ATML1* expression: one that occurs in P1 due to mechanical injury and another in P2 resulting from cell death in P1. Future studies to identify the cis-elements involved in *ATML1* expression specifically at P1 and P2, and forward and reverse genetic screens to uncover upstream regulators will be crucial to understanding how ATML1 exerts its distinct functions. Additionally, elucidating the different interacting partners for in ATML1 in P1 and P2 cells will enhance our understanding of how ATML1 performs distinct roles depending on the cell type.

Understanding wound healing in mature leaves will advance our knowledge of plant defense mechanisms and lay the groundwork for investigating plant–microbe interactions in this context. Additionally, as wounds trigger regeneration (Hoermayer et al., 2020; Ikeuchi et al., 2019; Iwase et al., 2021; Marhava et al., 2019; Yang et al., 2024; Zhou et al., 2019), a deep understanding of the wound-healing process will be essential for deciphering how the pathways to regeneration and recovery are determined. Moreover, our findings position the wound-healing process as a valuable model for exploring cell-type transitions and the associated hormonal regulatory networks, as well as gaining insights into the still mysterious positional cues of plant cells.

## ACKNOWLEDGMENTS

We thank N. Geldner, M.C. Seo, J.-H. Jung, D. Gasperini and Y. Joo for the generous sharing plant materials. We also thank Life Science Editors for manuscript editing. This work was supported by the Suh Kyungbae Foundation (SUHF-19010003) and the National Research Foundation of Korea (NRF-2021R1A5A1032428 and NRF-2023R1A2C1003077). K.K. was supported by the National Research Foundation of Korea (NRF-2022R1A6A3A0108604811) and J.-M.L. was supported by Hyundai Motor Chung Mong-Koo scholarship. J.-M.L. and M.H. were supported by the Stadelmann–Lee Scholarship Fund at Seoul National University, Korea.

## AUTHOR CONTRIBUTIONS

Y.L. and J.-M.L. conceived the study and designed the experiments. The experiments were performed by J.-M.L., M.H., M.K., K.K., S.J., G.H. and Y.Y. The computational analyses were performed by W.-T.J. and H.J. Y.L. and J.-M.L. drafted the manuscript, and all authors contributed to its revision. All authors have read and approved the final version.

## CONTACT FOR REAGENT AND RESOURCE SHARING

Further information and requests for resources and reagents should be directed to and will be fulfilled by the Lead Contact, Yuree Lee.

## SUPPLEMENTAL FIGURE LEGENDS

**Figure S1.**
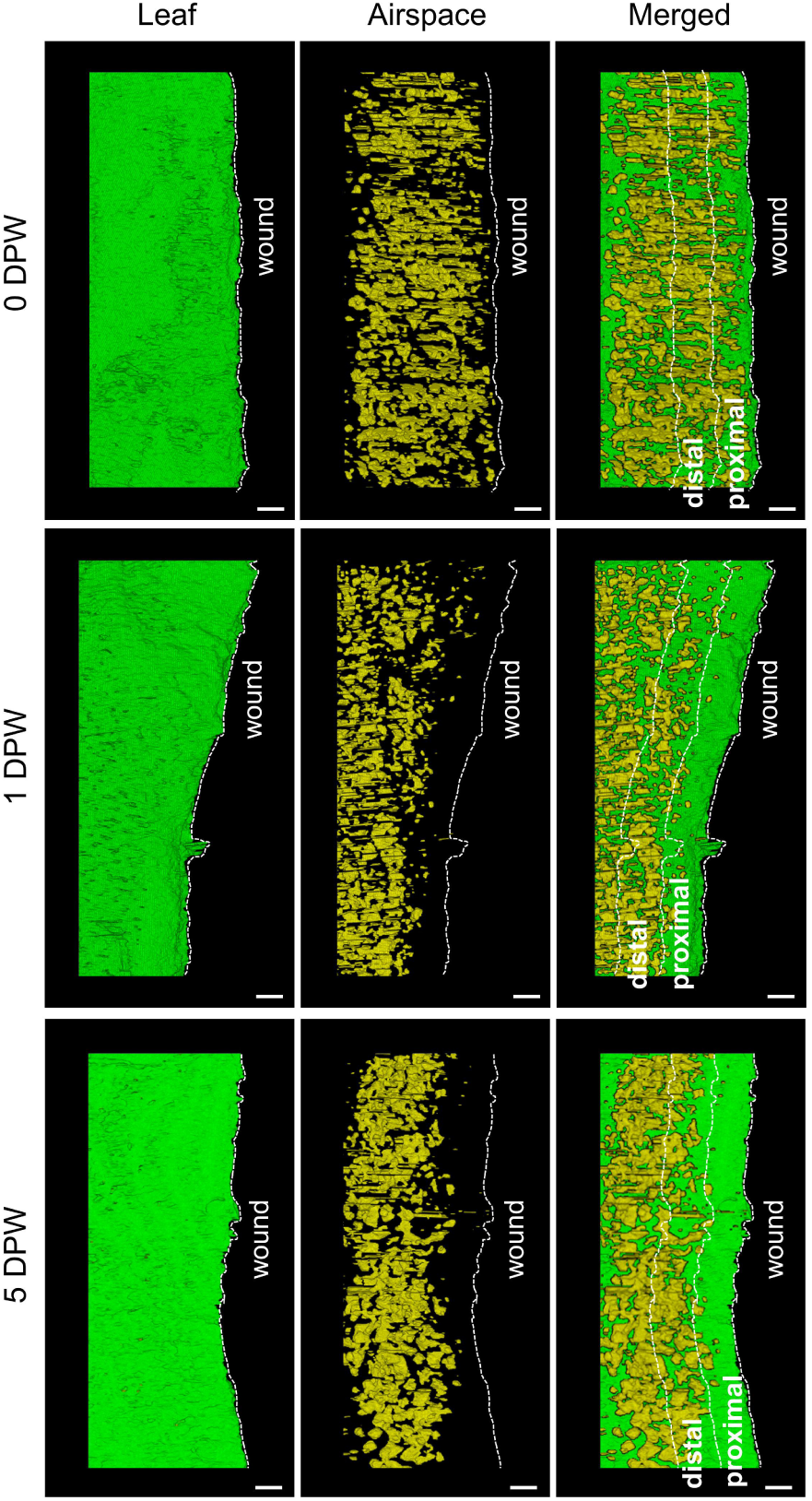
Three-dimensional cellular architecture of the wounded leaf, related to Figure 1. Maximum projection images of the wounded leaf were obtained using micro-CT at 0, 1, and 5 DPW. Scale bars, 100 μm.

**Figure S2.**
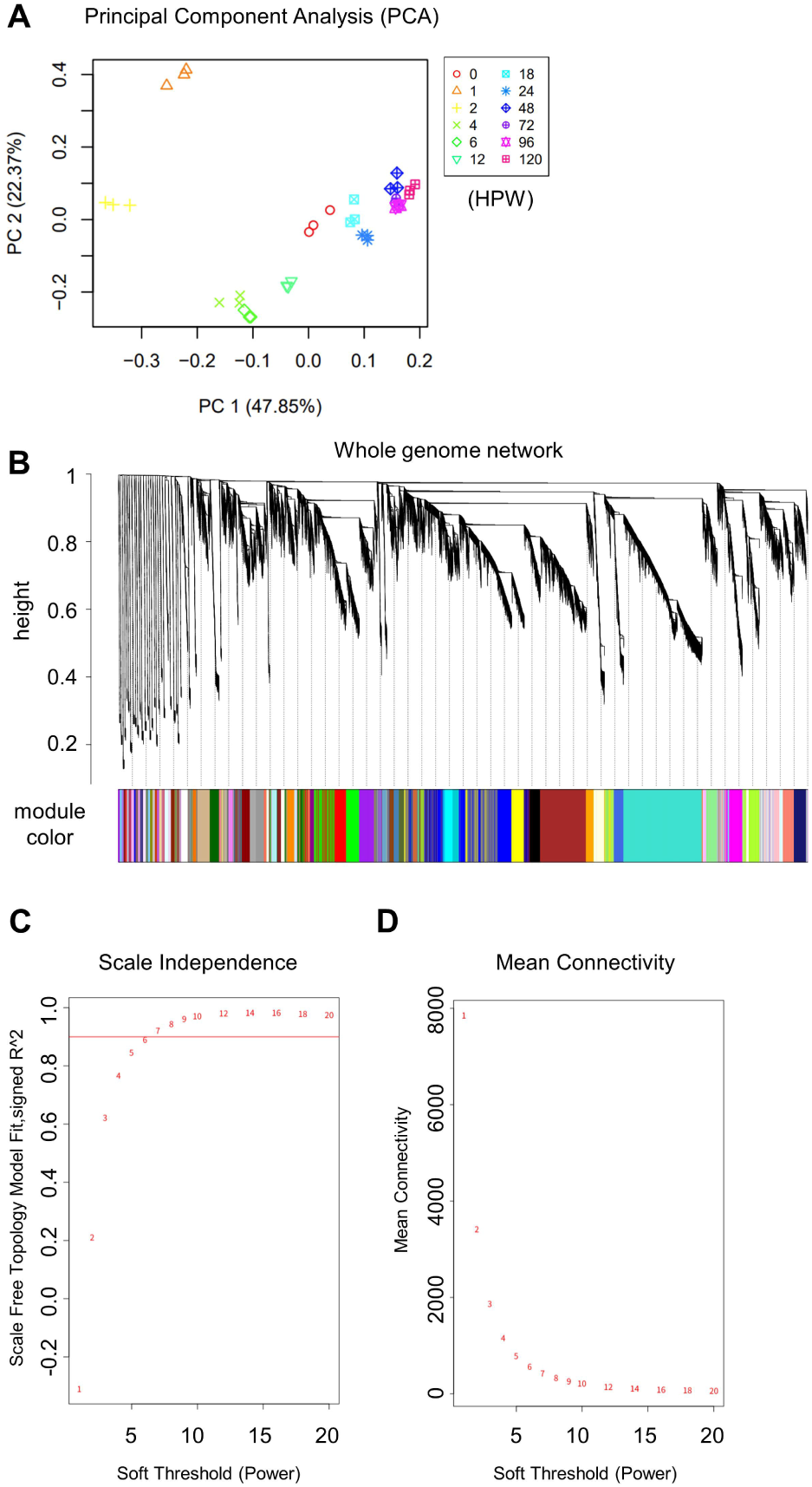
Overview of weighted gene co-expression network (WGCNA) analysis, related to Figure 2. (A) Principal Component Analysis (PCA) of all 36 time series samples. (B) WGCNA cluster dendrogram of all 36 time series samples, grouping genes into 23 distinct modules. (C and D) Scale independence and mean connectivity for calculating soft-thresholding powers.

**Figure S3.**
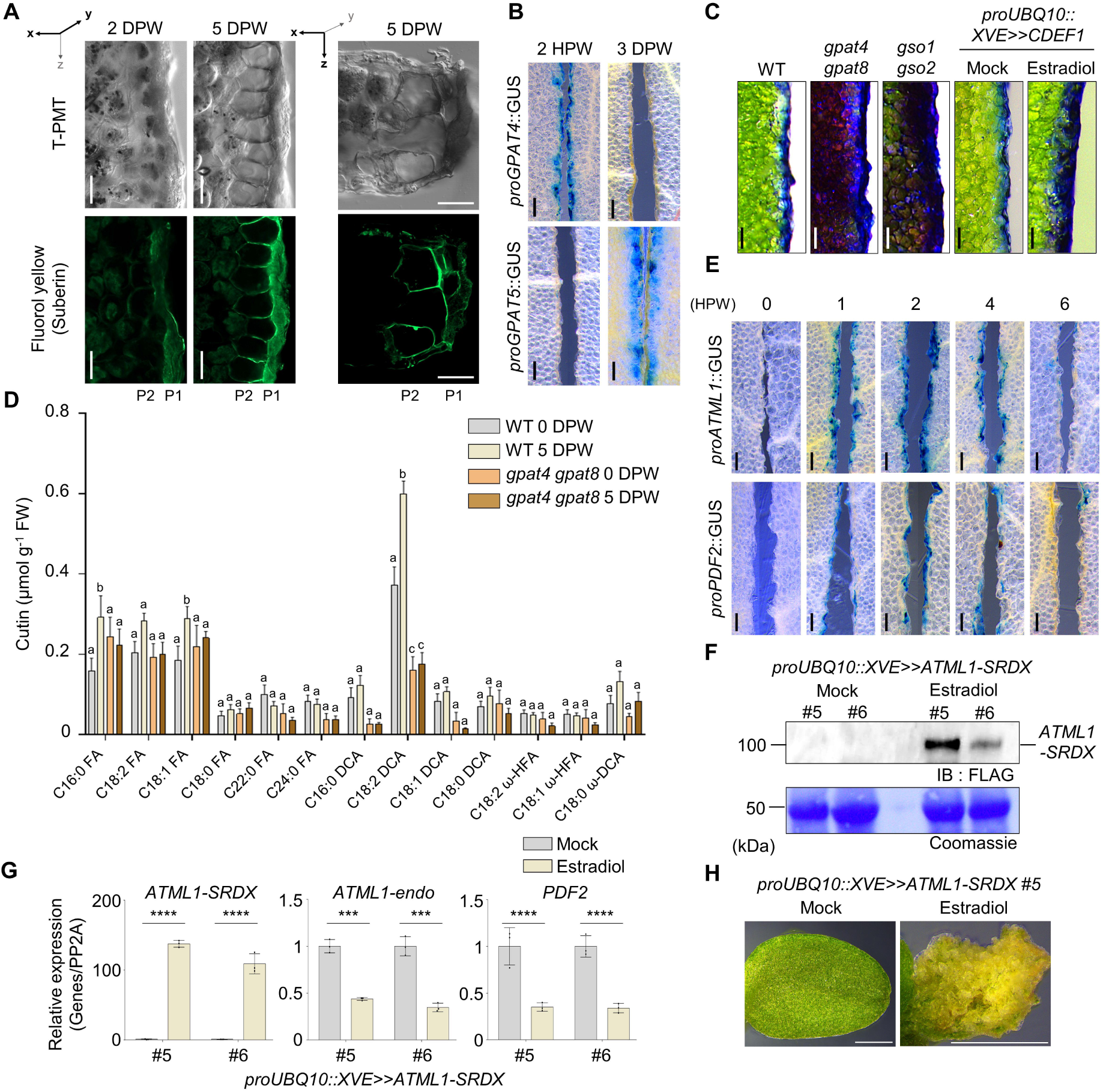
The wound surface is sealed by an ATML1-mediated cuticle at P1, related to Figure 3. (A) Confocal micrographs show suberin deposition in wounded leaves at 2 and 5 DPW, stained with Fluorol Yellow. The left two panels show palisade mesophyll cells from a top view, while the right panel presents a vertical cross-section of the leaf obtained with a vibratome. (B) Promoter-GUS analyses of *GPAT4* and *GPAT5* in the wounded site at 2 HPW and 3 DPW. (C) Toluidine blue (TB) staining shows the permeability of wounded site in WT and cuticle deficient mutants *gpat4 gpat8* and *gso1 gso2* and transgenic plants expressing *proUBQ10::XVE>>CDEF1* in the absence (mock) or presence of 10 μM estradiol. (D) Cutin composition in WT and *gpat4 gpat8* double-mutant wounded leaves at 0 HPW and 5 DPW. The error bars indicate mean ± SD. (E) Promoter-GUS histochemical analysis of *ATML1* and *PDF2* revealed that their expression at the wounded site began at 2 HPW. (F) Confirmation of *ATML1–SRDX* induction in transgenic plants expressing *proUBQ10::XVE>>ATML1-SRDX* using Western blot. Protein samples were extracted from leaf of 4-week-old *ATML1-SRDX* plants in the absence (mock) or presence of 10 μM estradiol for 1 day treatment. Mouse OctA-Probe Antibody was used to detect FLAG conjugated with SRDX, and coomassie blue was used to assess the Rubisco protein content, serving as a loading control for total protein. (G) Confirmation of *ATML1–SRDX* induction in transgenic plants expressing *proUBQ10::XVE>>ATML1-SRDX* using RT-qPCR. RNA was extracted from *ATML1-SRDX* plants in the absence (mock) or presence of 1 μM estradiol for 4 hour treatment. *ATML1-SRDX* expression was increased upon estradiol treatment. *ATML1-endo* and *PDF2* expression is down-regulated in estradiol treatment.—*N* = 3 biological replicates. *ATML1-SRDX*, reverse primer detects the SRDX sequence; *ATML1-endo*, reverse primer detects the 3′-untranslated region of ATML1 to examine *ATML1* endogeneous expression. (H) Representative images of deficient epidermal differentiation of 7-day-old *ATML1–SRDX* plants grown with 1 μM estradiol or Mock. In (D), statistical significance was determined by a one-sided Kruskal–Wallis test with Holm correction, followed by Dunn’s post-hoc test. Data points with different letters indicate significant differences representative of p < 0.05. In (G), statistical significance was determined by Student’s t-test (***p < 0.001, ****P < 0.0001). Scale bars, 50 μm (A), 100 μm (B, C, E), 500 μm (H).

**Figure S4.**
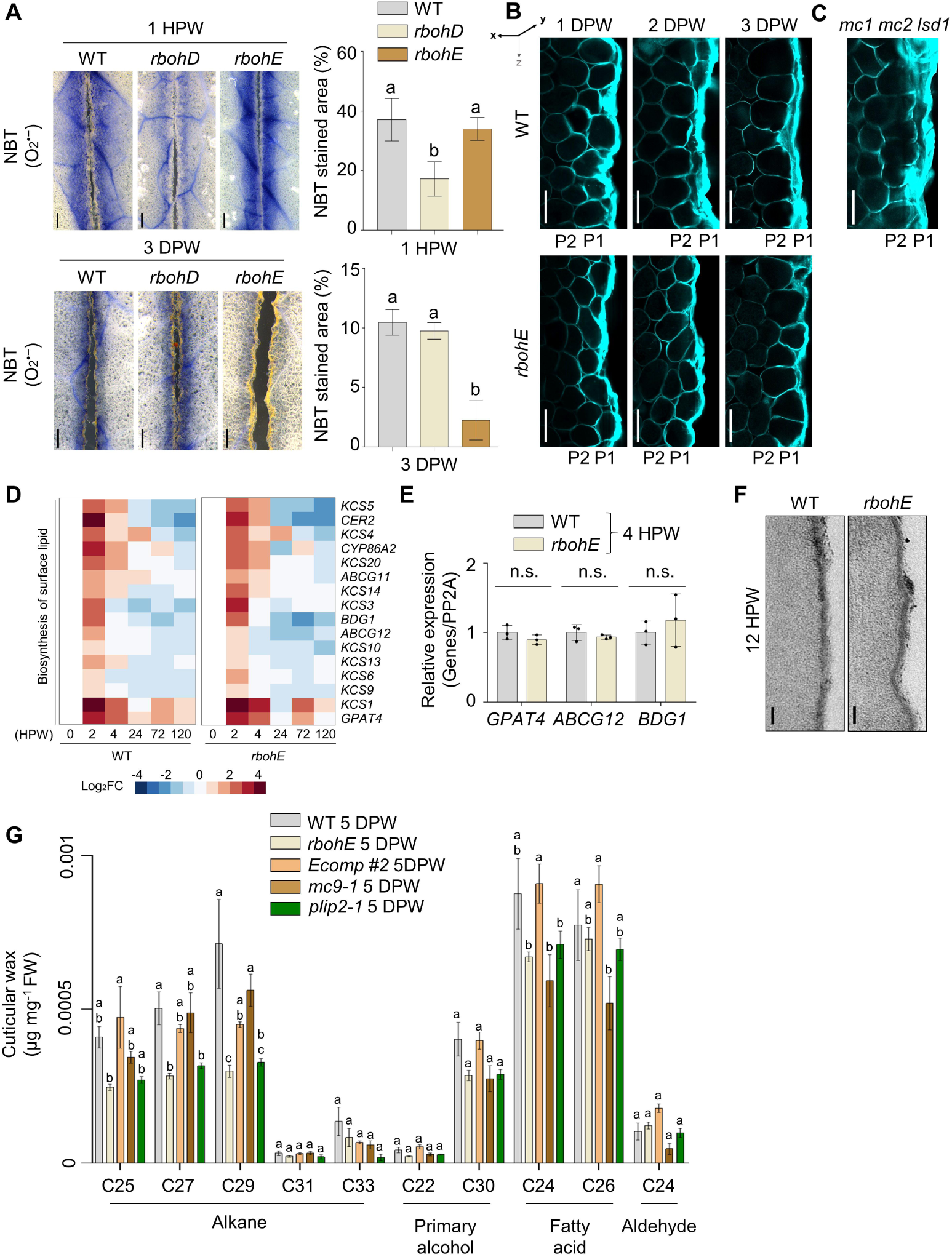
Ethylene–RbohE signaling is critical for PCD-mediated P1 maturation, related to Figure 4. (A) Superoxide accumulation detected with nitroblue tetrazolium (NBT) staining in wounded leaves of WT, *rbohD*, and *rbohE* at 1 HPW and 3 DPW. Quantification of the NBT stained area is shown (*n* = s10). (B and C) Confocal microscopy image of WT and *rbohE* at 1, 2, and 3 DPW (B) and *mc1 mc2 lsd1* at 5 DPW (C). Calcofluor-white was used to visualize the cell wall. (D) The heat map shows expression patterns of 2 - 4 HPW upregulated cutin and wax synthesis genes in WT and *rbohE*, and it is presented with a Log_2_FC scale relative to the 0-hour control for each background. See also Table S3. (E) Expression of the 2-4 HPW upregulated genes, *GPAT4, ABCG12,* and *BDG1* was analyzed by RT-qPCR in WT, *rbohE* at 4 HPW. The expression values were normalized to the PP2A gene, and calculated relative to the 0-hour control for each background. The expression levels are presented relative to the WT. The error bars indicate mean ± SD—*N* = 3 biological replicates. (F) Transmission electron micrographs of cuticle layer formation in WT and *rbohE* at 12 HPW. (G) Cuticular wax composition in wounded WT and various mutants at 5 DPW. The error bars indicate mean ± SD. In (A), statistical significance was analyzed using one-way ANOVA, followed by Tukey’s post-hoc test. In (E), statistical significance was determined by Student’s t-test (n.s. = no significance) Data points with different letters indicate significant differences representative of p < 0.05. Scale bars, 250 μm (A), 50 μm (B, C), 100 nm (F).

**Figure S5.**
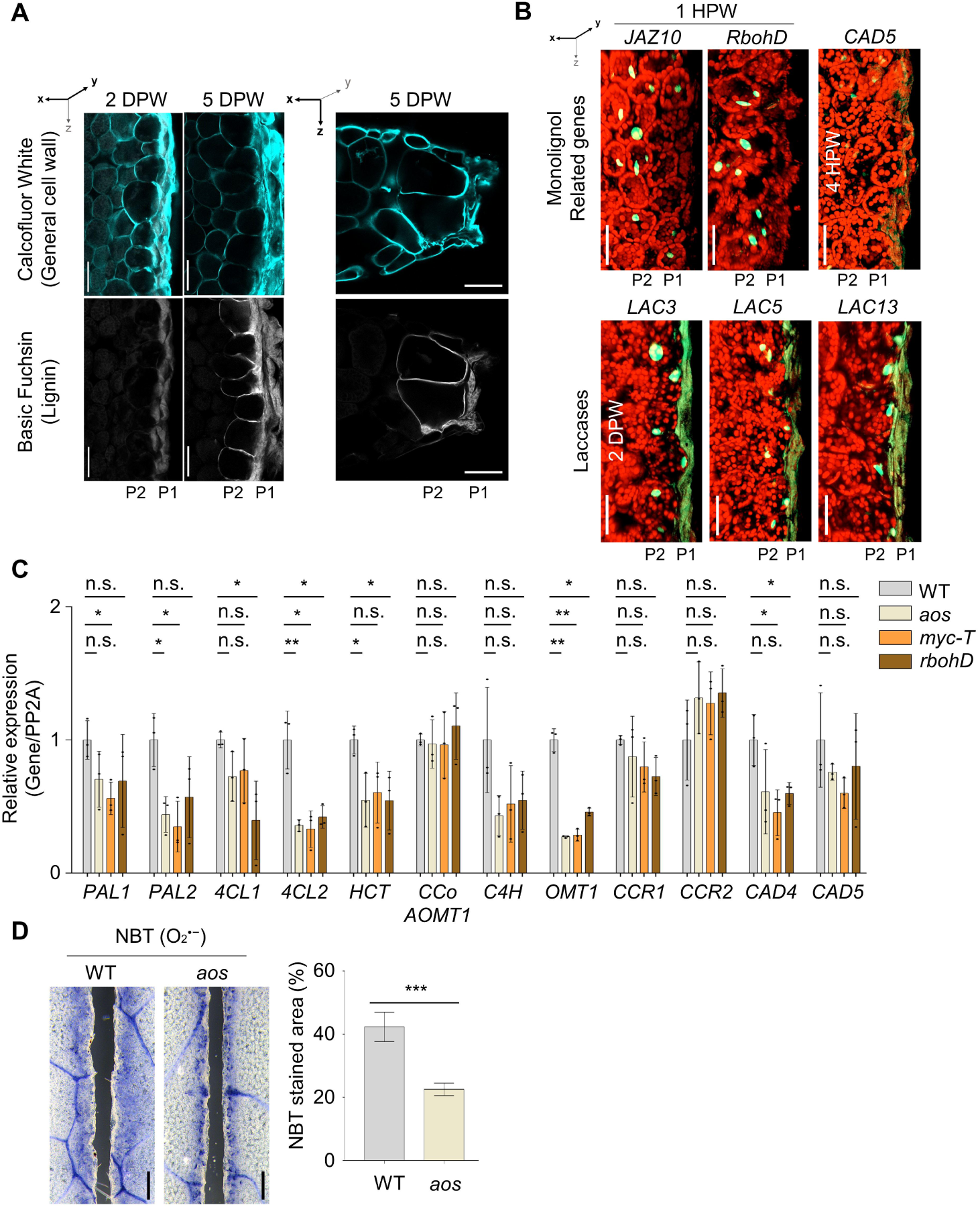
JA–RbohD signaling regulates the lignification of P2, related to Figure 5. (A) Confocal micrographs show lignin deposition of wounded leaves at 2 and 5 DPW. Basic fuchsin was used to stain lignin, and cell morphology was evaluated using calcofluor-white. The left two panels show palisade mesophyll cells from a top view, while the right panel presents a vertical cross-section of the leaf obtained with a vibratome. (B) Maximum projection images of confocal microscopy show the broad expression pattern of lignin synthesis-related genes and the P2-specific expression pattern of laccases. Green flourescence signal of *RbohD, CAD5, LAC3, LAC5,* and *LAC13* signals were visualized by promoter::nlsGFP and *JAZ10* by promoter::nls-3xVenus. (C) Expression of monolignol biosynthesis-related genes in WT and various mutants at 4 HPW. Expression levels were assessed using RT-qPCR, normalized to PP2A expression, and presented relative to WT. The error bars indicate mean ± SD—*N* = 3 biological replicates. (D) Superoxide accumulation detected with nitroblue tetrazolium (NBT) staining wounded leaves of WT and *aos* at 1 HPW and quantification of the NBT stained area (*n* = 10). In (C), statistical significance was determined by Student’s t-test (n.s. = no significance, *p < 0.05, **p < 0.01). In (D), statistical significance was analyzed using one-way ANOVA, followed by Tukey’s post-hoc test. Data points with different letters indicate significant differences representative of p < 0.05. Scale bars, 50 μm (A, B), 250 μm (D).

**Figure S6.**
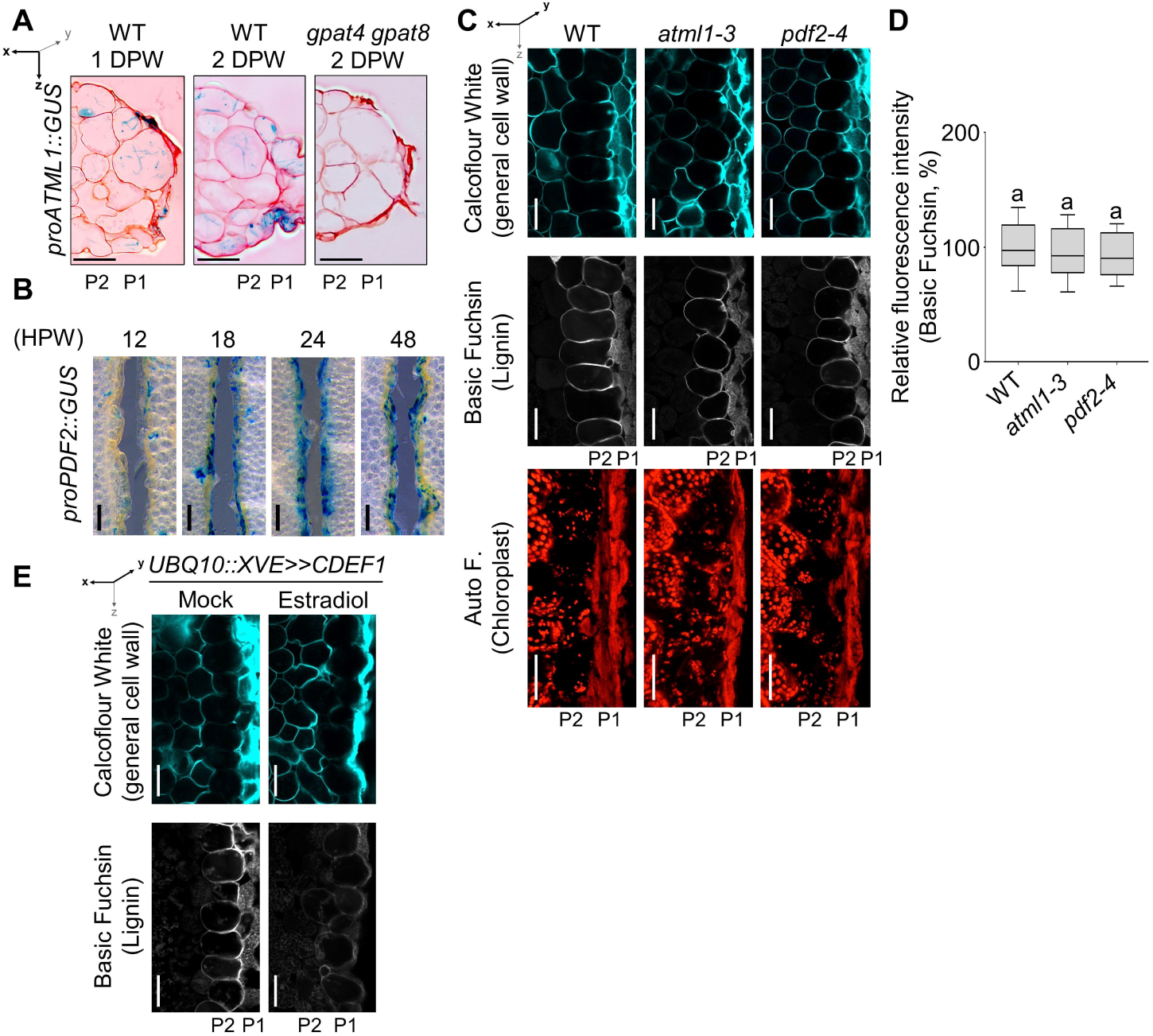
ATML1 module regulates P2 specification, related to Figure 6. (A) Section images of *ATML1* expression in WT and *gpat4 gpat8* mutants at 1 and 2 DPW. GUS stained cell was visualized with blue clour, and Sapranin staining was used to visualize cell morphology. (B) Promoter-GUS histochemical analyses of *PDF2* at 12 HPW to 2 DPW. (C and D) Wound-induced P2 specification in *atml1-3* and *pdf2-4* mutants. Lignification and chloroplast images were obtained from confocal microscopy (C) and lignin intensity quantification (D). The box and whisker plot shows data between the 10th and 90th percentiles. *n* = 100. (E) Confocal micrographs show P2 lignification in *UBQ10::XVE>>CDEF1* at 5 DPW. 10 μM or Mock treatment was applied to leaves before wounding. In (D), statistical significance was analyzed using one-way ANOVA, followed by Tukey’s post-hoc test. Data points with different letters indicate significant differences representative of p < 0.05. Scale bars, 50 μm (A, C, E), 100 μm (B).

## SUPPLEMENTAL TABLES

**Table S1.** GO analysis of WGCNA-clustered modules, related to Figure 2 (TMM sum>20; FDR < 0.05 by DAVID).

**Table S2.** TMM counts and Z-score of wound healing associated genes, related to Figure 2 and Figure 4 (TMM sum>20).

**Table S3.** List of differentially expressed genes related to surface lipid biosynthesis in WT vs. rbohE RNA-seq results, related to Figure 4.

**Table S4.** List of differentially expressed genes related to ethylene signaling and lignin polymerization in WT vs. gpat4 gpat8 RNA-seq results, related to Figure 6.

**Table S5.** List of primers used in this study, related to Star Methods.

## EXPERIMENTAL MODEL AND SUBJECT DETAILS

### *Arabidopsis thaliana* and growth conditions

Wild-type (WT) and mutant *Arabidopsis thaliana* possessed the Columbia (Col) genetic background. The following mutants were used in this study: *gpat4 gpat8* (Li et al., 2007), *gso1 gso2* (Tsuwamoto et al., 2008), *atml1-3* (Ogawa et al., 2015), *pdf2-4* (Ogawa et al., 2015), *rbohE* (Orman-Ligeza et al., 2016), *mc9-1* (Bollhoner et al., 2013), *etr1-1* (Gamble et al., 2002), *lsd1 atmc1 atmc2* (Coll et al., 2010), *wri4-1* (Park et al., 2016), *rbohD* (Torres et al., 2002), *aos* (Park et al., 2002), *myc2 myc3 myc4* (Wang et al., 2017), *9x lac (lac1;3;5;7;8;9;12;13;16)* (Rojas-Murcia et al., 2020), and *gpat5-1* (Beisson et al., 2007).

Arabidopsis seeds were surface sterilized and sown on half-strength Murashige and Skoog (MS) plates (4.4 g/L MS salt, 1 % (w/v) sucrose, pH 5.7, and 0.8% (w/v) agar), and stratified at 4 °C in the dark for 3 d. Then, the seedlings were transplanted in soil and grown in a growth chamber with 60% relative humidity under day-long conditions (16□h:8□h, dark:light cycles, respectively, 22 °C:18 °C, day:night regime, respectively, 70□μmol□m^−2^□s^−1^ photon flux density).

### Nicotiana benthamiana and growth conditions

*N. benthamiana* seeds were directly sown in the soil in 8 cm diameter pots. Seedlings were grown in a growth chamber and maintained at 60% relative humidity under day-long conditions (16□h:8□h, dark:light cycles, respectively, 22 °C:18 °C, day:night regime, respectively, 70□μmol□m^−2^□s^−1^ photon flux density). The first to the third fully expanded true leaves of 4-week-old plants were used for experiments.

### Capsella bursa-pastoris and growth conditions

*C. bursa-pastoris* seeds were harvested from Changnyeong, South Korea. The seeds used for the experiments were the third descendants of the field-collected seeds, all generations propagated by selfing. *C. bursa-pastoris* seeds were surface sterilized, sown on half-strength Murashige and Skoog (MS) plates (4.4 g/L MS salt, 1 % (w/v) sucrose, pH 5.7, and 0.8 % (w/v) agar), and stratified at 4 °C in the dark for 3 d. Then, the seedlings were transplanted in soil and grown in a growth chamber with 60% relative humidity under day-long conditions (16□h:8□h, dark:light cycles, respectively, 22 °C:18 °C, day:night regime, respectively, 70□μmol□m^−2^ s^−1^ photon flux density). 8^th^ leaf of 4-week-old plants were used for experiments.

## METHOD DETAILS

### Plasmid Constructions and Plant Transformations

Gateway Cloning Technology (Invitrogen) was used to generate plasmid constructs. The list of plasmid constructs and primer details are presented in Table S5. SRDX-3xFLAG construct was synthesized from Macrogen. The plasmid constructs were introduced into plants using the floral dipping method (Martinez-Trujillo et al., 2004). Briefly, Arabidopsis flowers were dipped into a solution containing Agrobacterium tumefaciens, 5% sucrose, and 0.05% surfactant Silwet L-77 for 2 to 3 s with gentle agitation. Dipped plants were maintained under high humidity conditions for 16 to 24 hours. Transformed plants were selected using antibiotics or herbicide selection markers.

### Pharmacological Assays

To analyze the drug effects, the drug was infiltrated into the leaves before wound induction. A total of 10 µM methyl jasmonic acid (MeJA, Sigma-Aldrich), 1–10 µM estradiol, 100 µM diphenyleneiodonium (DPI, Sigma-Aldrich), or 50 µM salicylhydroxamic acid (SHAM, Sigma-Aldrich) was used. For monolignol treatment, 30 µM coniferyl alcohol (Sigma-Aldrich) and 30 µM sinapyl alcohol (Sigma-Aldrich) were infiltrated in PBS.

### Wound Treatment and Subsequent Sampling for Further Analysis

The eighth leaves of 4-week-old Arabidopsis thaliana were mechanically wounded using stainless-steel blades (size 10, FEATHER) attached to a laboratory knife. The wounds were created with the ridges aligned parallel to the primary vasculature of the leaf, and care was taken to ensure vertical incisions to observe the P1 and P2 morphologies. Then, the wounded plants were transferred to a growth chamber and incubated for the required duration. To standardize the plant wound response, the initiation point of the wound was carefully fixed, and plants from the same tray were used.

For leaf sampling intended for imaging, the samples were fixed in 4% (w/v) paraformaldehyde at room temperature for 1 hour before being briefly washed with 1x PBS. Next, the samples were transferred to ClearSee solution until the chlorophyll was completely bleached (Ursache et al., 2018).

For leaf sampling intended for RNA extraction, samples were quickly excised using stainless-steel blades (size 10, FEATHER) to a depth of 5 mm from the wounded site and immediately placed in 2 mL Eppendorf tubes submerged in liquid nitrogen to prevent the upregulation of early wound signals induced by mechanical damage during sampling. For reliable experimental results, a minimum of 8 individual leaf samples were included for each biological replicate.

For leaf sampling intended for lipid extraction, samples were quickly excised using stainless-steel blades (size 10, FEATHER) to a depth of 5 mm from the wounded site, and their weights were measured. The samples were immediately transferred to glass extraction tubes and incubated in liquid nitrogen. For each biological sample, 0.5 mg of tissue was recommended, which typically requires at least 15 individual plants. For each genotype, a minimum of four independent replicates were performed in a single experiment.

### Microscopy and Visualization

Low-magnification images were captured using a Leica M205FA stereo microscope (Leica). Confocal laser scanning microscopy was performed using an LSM 900 confocal microscope (Zeiss) with the following excitation and detection setups: GFP: 488 nm, 500– 545 nm; mCherry: 594 nm, 570–670 nm; calcofluor-white: 405 nm, 430–480 nm; basic fuchsin: 561 nm, 580–670 nm; Fluoral Yellow: 405 nm, 430–500 nm.

Cleared samples were stained using 0.4 % basic fuchsin (Sigma-Aldrich) in Clearsee solution at room temperature overnight to visualize the lignin structure. Then, samples were washed twice in Clearsee solution for at least 1 hour each time. Next, samples were transferred in 0.2% calcofluor-white (Sigma-Aldrich) in Clearsee solution and incubated overnight (Ursache et al., 2018). Fluorescence images were obtained using an LSM 900 confocal microscope (Zeiss).

To visualize the suberin structure, samples were incubated in a freshly prepared solution of Fluoral Yellow 088 (Santa Cruz biotechnology) (0.01% w/v, in lactic acid) at 70 °C for 90 min. Then, the samples were washed in water for 30 min. Lastly, each sample was mounted using a 50% glycerol solution. Fluorescence images were obtained using an LSM 900 confocal microscope (Zeiss).

To visualize superoxide accumulation, leaves were stained using nitroblue tetrazolium (NBT, Sigma-Aldrich) (Straus et al., 2010). Briefly, the leaves were vacuum-infiltrated using a NBT solution (0.05% in PBS) and incubated in the dark for 2 hours. After NBT staining, the leaves were fixed and destained in 3:1:1 ethanol:lactic acid:glycerol (bleaching solution) at 70 °C until chlorophyll was completely removed. Subsequently, each leaf was visualized using a Leica M205FA stereo microscope.

For the permeability assay, leaves were treated with toluidine blue O (Fisher Scientific) solution in water (0.025%) for 10 min and washed with water for 5 min. Images were obtained using a Leica M205FA stereo microscope.

For β-glucuronidase (GUS) staining, leaves were fixed in 90% acetone (Dedow et al., 2022) and incubated in GUS solution (3□mM potassium ferrocyanide, 3□mM potassium ferricyanide, 1□mM X-Gluc, 1□M NaH_2_PO_4_, 1□M Na_2_HPO_4_, 0.5□M EDTA, pH□8.0) at 37 °C for 24 h. After staining, leaves were incubated in 70% (v/v) ethanol until chlorophyll was completely removed and then visualized using a Leica M205FA stereo microscope.

### Histology

GUS-stained and non-stained leaves were sectioned to visualize the vertical structure. For wound-induced GUS histology, samples were fixed in 90% acetone for 20 minutes under vacuum and then at room temperature for 1 hour (Dedow et al., 2022). The samples were processed for histological analysis.

For resin-embedded sections, leaves were fixed in 4% paraformaldehyde (PFA) for 1 hour, followed by a dehydration series using ethanol concentrations ranging from 30% to 100%. Subsequently, the samples were pre-infiltrated with stepwise dilutions of Technovit 7100 solution (Kluzer Technik) in ethanol at different ratios (1:5, 2:5, 3:5, 4:5, 5:5, 3.5:2.5, 4.5:2.5, 5.5:2.5, 6.5:2.5, 7.5:2.5, v/v) for 1 hour each. The resulting resin blocks were then sectioned into 4 μm slices using a Leica Histocore AUTOCUT rotary microtome (Leica) and stained with either 0.01% (w/v) toluidine blue (ThermoFisher Scientific) or 0.05% (w/v) safranin (Sigma-Aldrich). For paraffin-embedded sections, leaves were fixed in 4% PFA for 1 hour and then dehydrated through a graded ethanol series (30% to 100%). Following dehydration, the samples were cleared with xylene and infiltrated with paraffin wax at 60 °C. The paraffin-embedded samples were then sectioned into 5 μm slices using a rotary microtome, and the sections were mounted onto glass slides. The slides were then dewaxed in xylene, rehydrated through a graded ethanol series, and used for TUNEL assay.

### Scanning and Transmission Electron Microscopy

For scanning electron microscopy (SEM), tissue samples were first subjected to critical point drying using an SPI DRY critical point dryer (SPI Supplies) to preserve their structural integrity. Following drying, the samples were mounted onto steel stubs using conductive carbon adhesive. The mounted samples were then coated with a thin layer of osmium using an osmium coater to enhance conductivity and improve image resolution. The coated specimens were subsequently examined using a JSM 6390LV scanning electron microscope (JEOL) under high-vacuum conditions.

For transmission electron microscopy, leaves were fixed at 4 °C overnight using 2% paraformaldehyde and 2% glutaraldehyde in 50 mM potassium phosphate buffer (pH 7.0), which was followed by post-fixation with 1% w/v osmium tetroxide at 4 °C for 2 h. The samples were washed three times in distilled water and dehydrated through a series of ethanol incubations (30%, 50%, 70%, 80%, and 90% for 30 min at each step) and 100% ethanol (30 min; 3 times). The samples were then infiltrated with Spurr’s resin (EMS) and polymerized at 70 °C for 48 h. Ultrathin sections of 50 nm thickness were analyzed using a Tecnai G2 Spirit transmission electron microscope (FEI, ThermoFisher Scientific, SNU CMCI) at 120 kV.

### X-ray Micro-computed Tomography

X-ray micro-computed tomography (micro-CT) of leaves was conducted following the previously described procedure with slight modifications (Kim et al., 2024). Leaves were placed inside 1.5 mL Eppendorf tubes coated with a thin layer of lip balm (Nivea) to prevent movement, and 20 μL of distilled water was added to maintain hydration. To visualize the anatomical structure of the leaves, micro-CT scans were performed using a Skyscan 1272 system (Bruker) at a resolution of 3 μm, with a field of view of 2016 × 2016 pixels. Scans were conducted using an electron acceleration energy of 50 kV, a current of 200 μA, and a detector exposure time of 700 ms. The raw image data were reconstructed using NRecon software (version 1.7.4.2, Bruker) and reoriented using DataViewer software (Ordinola-Zapata et al., 2017). Quantification of leaf tissue and porosity, defined as the proportion of airspace relative to the total volume, was performed using CTAn (version 1.18, Bruker) in accordance with the following guidelines provided by Bruker Micro-CT. The final 3D structures were rendered using CTVox (Version 3.3, Bruker) software (Liu et al., 2018), and color-coded images representing airspace calculated as the diameter of the largest sphere that can fit within a pore were generated using CTAn (Kim et al., 2024).

### Cell Death Staining

Apoptotic cells in paraffin-embedded sections were identified using the DeadEnd Fluorometric TUNEL (terminal deoxynucleotidyl transferase dUTP nick end labeling) system (Promega) (Tripathi et al., 2017). Transverse sections were subjected to TUNEL staining following the manufacturer’s instructions and subsequently mounted using Vectashield antifade mounting medium with DAPI (Vector Laboratories) for visualization. Fluorescence signals were captured by a LSM 900 confocal microscope (Zeiss), utilizing an excitation wavelength of 488 nm and emission detection between 500 and 540 nm for TUNEL, and an excitation wavelength of 358 nm with emission detection between 440 and 490 nm for DAPI.

### Western Blotting

Leaves of transgenic plants overexpressing *ATML1-SRDX* infiltrated with either 10 μM estradiol (Sigma-Aldrich) or DMSO (Sigma-Aldrich) were harvested and homogenized using a TissueLyser II (QIAGEN) in the presence of liquid nitrogen. Total protein was extracted by resuspending the ground tissue in a lysis buffer composed of 10 mM Tris–HCl (pH 7.5), 150 mM NaCl, 0.5 mM EDTA, 0.5% (v/v) NP-40, and 0.09% (w/v) sodium azide. Next, the extract was centrifuged at 13,000 g and 4 °C for 20 minutes to remove debris. The supernatant containing the total protein was subjected to SDS-PAGE, utilizing a 12.5% separating gel and a 4% stacking gel. Following electrophoresis, proteins were transferred to a membrane, and immunoblotting was performed. After blocking the membrane in 5% (w/v) low-fat dry milk for 2 hours, the membrane was then incubated with an Mouse OctA-Probe Antibody (H-5) (Santa Cruz Biotechnology) at a dilution of 1:2000 in Tris-buffered saline containing 0.1% (v/v) Tween-20 (TBST) at room temperature for 2 hours. The ATML1-SRDX protein was detected using a Fusion-solo ChemiDoc (VILBER). Coomassie blue staining (Sigma-Aldrich) was applied to the membrane to assess the Rubisco protein content, serving as a loading control for total protein.

### Cutin Analysis

Cutin analysis was conducted on wounded leaves following a modified version of a previously described protocol (Lee et al., 2021). Briefly, samples were collected from leaf tissue approximately 5 mm from the wound site, and the fresh weight of 15 individual leaves was measured. The leaves were then immersed in hot isopropanol for 10 minutes. Hexane was added in a 2:3 ratio into isopropanol, and the samples were subjected to delipidation using a series of chloroform mixtures (2:1, 1:1, and 1:2, v/v) over two days, followed by treatment with pure methanol for 30 minutes, until the tissue appeared colorless. The delipidated tissues were then depolymerized via sulfuric acid-catalyzed reaction at 85 °C for 2 hours in the presence of toluene, using methyl heptadecanoate (Sigma-Aldrich) and ω-pentadecalactone (Sigma-Aldrich) as internal standards. After cooling to room temperature, 1.5 mL of 0.9% (w/v) KCl solution and 1 mL of dichloromethane were added. The organic phase was evaporated to dryness under nitrogen (N_2_) gas, and the samples were derivatized by adding 100 μL of pyridine (Sigma-Aldrich), incubating at 60 °C for 1 hour. The acetylated samples were evaporated under nitrogen gas and dissolved in a heptane (1:1, v/v) mixture for gas chromatography–mass flame ionization detector (GC–FID, GC-2030, Shimadzu, Japan) analysis; for GC–FID, a DB-5 capillary column (30 m, 0.25 mm inner diameter, 0.25 μm film thickness; J/W Scientific) was employed.

### Cuticular Wax Analysis

Cuticular wax analysis was performed on 15 wounded leaf samples using a previously established method with some modifications (Lee et al., 2021). In brief, the cuticular waxes were extracted by shaking the leaf samples in chloroform for 30 seconds. The chloroform extracts were then evaporated under nitrogen (N_2_) gas and redissolved in a solvent mixture of N,O-bis(trimethylsilyl) trifluoroacetamide (BSTFA, Sigma-Aldrich) and pyridine (Sigma-Aldrich) (1:1, v/v). The samples were incubated at 90 °C for 30 minutes. Following incubation, the samples were centrifuged at 110 g and 25 °C for 5 minutes. The silylated samples were then evaporated under N_2_ gas and reconstituted in a heptane/toluene mixture (1:1, v/v). The composition and quantity of the cuticular waxes were analyzed using GC–MS with a QP2010 system and gas chromatography (GC) with a GC-2010 system (GCMS-QP2010 SE, Shimadzu).

### RNA Extraction, cDNA Synthesis, and RT-qPCR

Total RNA was extracted from the wounded leaf samples using the RNeasy Plant Mini kit (Qiagen) following the manufacturer’s protocol. RNA concentration and purity were assessed using an ND-2000 spectrophotometer (ThermoFisher Scientific), with RNA samples showing optical density (OD) 260/280 ratios between 1.9 and 2.1 and OD 260/230 ratios between 2.0 and 2.5. RNA integrity was further evaluated using a TapeStation RNA screen tape (Agilent, Santa Clara, CA, USA), and only samples with an RNA integrity number (RIN) greater than 8.0 were selected for RNA sequencing (RNA-Seq) library preparation.

For complementary DNA (cDNA) synthesis, 1 μg of total RNA was reverse-transcribed using an amfiRivert cDNA Synthesis mix (GenDepot) with oligo (dT) primers, according to the manufacturer’s instructions. The resulting cDNA was then used for quantitative polymerase chain reaction (qPCR) analysis on a Quant Studio 1 system (Applied Biosystems) with SYBR Green Real-time PCR Master Mix (ThermoFisher Scientific). Primer sequences are provided in Supplementary Table S5. Gene expression levels were calculated using the threshold cycle (Ct) method, and the relative expression of target genes was determined using the ΔΔCt method: ΔΔCt = (Ct target gene – Ct PP2A gene) in treated samples – (Ct target gene – Ct PP2A gene) in control samples (Livak and Schmittgen, 2001).

### RNA Sequencing

Total RNA was extracted from wounded leaves of *A.thaliana* using a Qiagen RNeasy Mini kit (Qiagen) per the manufacturer’s instructions. Samples with an RNA integrity number greater than 0.8 were selected for library construction. After assessing the quality and quantifying the RNA, mRNA-sequencing libraries were prepared using the TruSeq Stranded mRNA Library Prep kit (Illumina) according to the provided protocol. The adaptor-ligated libraries were then sequenced using the Illumina HiSeq 2500 platform (Illumina).

The TruSeq universal and indexed adapters were trimmed using Trimmomatic software v0.39 (Bolger et al., 2014). The reads obtained from sequencing were filtered using PRINSEQ v0.20.4 (Schmieder and Edwards, 2011). The filtered reads were aligned to the TAIR10.1 reference genome (GCF_000001735.4) using HISAT2 v2.2.1 (Kim et al., 2019). Gene abundances were quantified with StringTie v1.3.4 (Pertea et al., 2015), utilizing the TAIR10 genome GFF3 annotation file for reference gene identification. The resulting gene-level expression data were compiled into a matrix representing expression levels in transcripts per million (TPM). To account for compositional biases across samples, we applied trimmed mean of M-values (TMM) normalization using the edgeR package v3.19 in R (Robinson et al., 2010). Genes for which the sum of the TMM means from each of the three duplicates was less than 20 were filtered out. Finally, z-scores or log2 fold change values for the target gene list were visualized as a heatmap using the pheatmap package v1.0.12 in R (Kolde, 2019).

### WGCNA and Gene Ontology

A signed co-expression network was constructed utilizing the WGCNA v1.72 package in R, incorporating all 36-time series samples (Langfelder and Horvath, 2008). The network construction commenced with the computation of pairwise correlations between genes across the samples, followed by generating an adjacency matrix by raising the co-expression values (0.5□+□0.5□×□correlation matrix) to the power of β = 6, serving as a soft threshold. The adjacency matrix was converted into a topological overlap matrix (TOM), providing a robust an d biologically relevant measure of network interconnectedness. Subsequent average linkage hierarchical clustering facilitated grouping genes with similar co-expression patterns. Gene modules were delineated as branches using the Dynamic Tree Cut algorithm applied to the hierarchical clustering dendrogram. Gene lists within each co-expression module were also subjected to Gene Ontology (GO) analysis using the DAVID platform (Huang da et al., 2009).

### Quantification and Statistical Analysis

Lignin and suberin intensity were measured using Image J software with the following formula: i) region of interest (ROI) was established to quantify fluorescence. ii) Mean fluorescence was calculated from 5 ROIs for single cell results. The fluorescence intensity of the inhibitor-treated samples or mutants was compared with the untreated control or WT. Figure 5C-D, F-G, H-I, and K-L show the relative values (%) as the mean (n = 100 cells) ± SD. These data were subjected to ANOVA along with the one-way ANOVA, followed by post hoc Tukey’s test for significance.

Toluidine blue and cuticle thickness were measured using Image J software within the following formula: i) 10 individual linear depths were measured. ii) Mean depth was calculated from 10 individual data points. The depth was measured from 5 individual leaves, which showed linear results from 3 independent experiments in Figure 1A-B, 3E-F, 4E, and 5A. The cuticle thickness was measured from 5 individual leaves from 3 independent experiments in Figure 4G. These data were subjected to one-way ANOVA, followed by post hoc Tukey’s test for significance. The statistical analyses for all experiments were performed with the Graphpad prism 10.0 software (https://www.graphpad.com/). The number of biological replicates and the statistical details of each experiment were described in the related figure legends.

### Data availability statement

The raw data files for the RNA-seq analysis reported in this paper can be found at GenBank under the accession number GSE275743.

## REFERENCES

Abe, M., Katsumata, H., Komeda, Y., and Takahashi, T. (2003). Regulation of shoot epidermal cell differentiation by a pair of homeodomain proteins in Arabidopsis. Development 130, 635–643.

Arya, G.C., Sarkar, S., Manasherova, E., Aharoni, A., and Cohen, H. (2021). The Plant Cuticle: An Ancient Guardian Barrier Set Against Long-Standing Rivals. Front Plant Sci 12, 663165.

Barros, J., Serk, H., Granlund, I., and Pesquet, E. (2015). The cell biology of lignification in higher plants. Ann Bot 115, 1053–1074.

Barton, K.A., Schattat, M.H., Jakob, T., Hause, G., Wilhelm, C., McKenna, J.F., Mathe, C., Runions, J., Van Damme, D., and Mathur, J. (2016). Epidermal Pavement Cells of Arabidopsis Have Chloroplasts. Plant Physiol 171, 723–726.

Beaudoin, F., Wu, X., Li, F., Haslam, R.P., Markham, J.E., Zheng, H., Napier, J.A., and Kunst, L. (2009). Functional characterization of the Arabidopsis beta-ketoacyl-coenzyme A reductase candidates of the fatty acid elongase. Plant Physiol 150, 1174–1191.

Beisson, F., Li, Y.H., Bonaventure, G., Pollard, M., and Ohlrogge, J.B. (2007). The acyltransferase GPAT5 is required for the synthesis of suberin in seed coat and root of. Plant Cell 19, 351–368.

Berhin, A., de Bellis, D., Franke, R.B., Buono, R.A., Nowack, M.K., and Nawrath, C. (2019). The Root Cap Cuticle: A Cell Wall Structure for Seedling Establishment and Lateral Root Formation. Cell 176, 1367-+.

Bolger, A.M., Lohse, M., and Usadel, B. (2014). Trimmomatic: a flexible trimmer for Illumina sequence data. Bioinformatics 30, 2114–2120.

Bollhoner, B., Zhang, B., Stael, S., Denance, N., Overmyer, K., Goffner, D., Van Breusegem, F., and Tuominen, H. (2013). Post mortem function of AtMC9 in xylem vessel elements. New Phytol 200, 498–510.

Canher, B., Heyman, J., Savina, M., Devendran, A., Eekhout, T., Vercauteren, I., Prinsen, E., Matosevich, R., Xu, J., Mironova, V., et al. (2020). Rocks in the auxin stream: Wound-induced auxin accumulation and ERF115 expression synergistically drive stem cell regeneration. Proc Natl Acad Sci U S A 1 17, 16667–16677.

Christiaens, F., Canher, B., Lanssens, F., Bisht, A., Stael, S., De Veylder, L., and Heyman, J. (2021). Pars Pro Toto: Every Single Cell Matters. Front Plant Sci 12, 656825.

Coen, O., Lu, J., Xu, W.J., De Vos, D., Pechoux, C., Domergue, F., Grain, D., Lepiniec, L., and Magnani, E. (2019). Deposition of a cutin apoplastic barrier separating seed maternal and zygotic tissues. B mc Plant Biol 19.

Coll, N.S., Vercammen, D., Smidler, A., Clover, C., Van Breusegem, F., Dangl, J.L., and Epple, P. (2010). Arabidopsis type I metacaspases control cell death. Science 330, 1393–1397.

De Bellis, D., Kalmbach, L., Marhavy, P., Daraspe, J., Geldner, N., and Barberon, M. (2022). Extracellular vesiculo-tubular structures associated with suberin deposition in plant cell walls. Nat Commun 13, 1489.

Dedow, L.K., Oren, E., and Braybrook, S.A. (2022). Fake news blues: A GUS staining protocol to reduce false-negative data. Plant Direct 6, e367.

Delude, C., Moussu, S., Joubes, J., Ingram, G., and Domergue, F. (2016). Plant Surface Lipids and Epidermis Development. Subcell Biochem 86, 287–313.

Denness, L., McKenna, J.F., Segonzac, C., Wormit, A., Madhou, P., Bennett, M., Mansfield, J., Zipfel, C., and Hamann, T. (2011). Cell wall damage-induced lignin biosynthesis is regulated by a reactive oxygen species-and jasmonic acid-dependent process in Arabidopsis. Plant physiology 156, 1364–1374.

Dixon, R.A., and Barros, J. (2019). Lignin biosynthesis: old roads revisited and new roads explored. Open Biol 9, 190215.

Espelie, K.E., Sadek, N.Z., and Kolattukudy, P.E. (1980). Composition of suberin-associated waxes from the subterranean storage organs of seven plants : Parsnip, carrot, rutabaga, turnip, red beet, sweet potato and potato. Planta 148, 468–476.

Fernandez-Calvo, P., Chini, A., Fernandez-Barbero, G., Chico, J.M., Gimenez-Ibanez, S., Geerinck, J., Eeckhout, D., Schweizer, F., Godoy, M., Franco-Zorrilla, J.M., et al. (2011). The Arabidopsis bHLH transcription factors MYC3 and MYC4 are targets of JAZ repressors and act additively with MYC2 in the activation of jasmonate responses. Plant Cell 23, 701–715.

Gamble, R.L., Qu, X., and Schaller, G.E. (2002). Mutational analysis of the ethylene receptor ETR1. Role of the histidine kinase domain in dominant ethylene insensitivity. Plant Physiol 128, 1428–1438.

Gechev, T.S., Van Breusegem, F., Stone, J.M., Denev, I., and Laloi, C. (2006). Reactive oxygen species as signals that modulate plant stress responses and programmed cell death. Bioessays 28, 1091–1101.

Gorczyca, W., Gong, J., and Darzynkiewicz, Z. (1993). Detection of DNA strand breaks in individual a poptotic cells by the insitu terminal deoxynucleotidyl transferase and nick translation assays. Cancer research 53, 1945–1951.

Graca, J., and Santos, S. (2006). Glycerol-derived ester oligomers from cork suberin. Chem Phys Lipids 144, 96–107.

Hoermayer, L., Montesinos, J.C., Marhava, P., Benkova, E., Yoshida, S., and Friml, J. (2020). Wounding-induced changes in cellular pressure and localized auxin signalling spatially coordinate restorative divisions in roots. Proc Natl Acad Sci U S A 117, 15322–15331.

Hoermayer, L., Montesinos, J.C., Trozzi, N., Spona, L., Yoshida, S., Marhava, P., Caballero-Mancebo, S., Benkova, E., Heisenberg, C.P., Dagdas, Y., et al. (2024). Mechanical forces in plant tissue matrix orient cell divisions via microtubule stabilization. Dev Cell 59, 1333–1344 e1334.

Huang da, W., Sherman, B.T., and Lempicki, R.A. (2009). Bioinformatics enrichment tools: paths toward the comprehensive functional analysis of large gene lists. Nucleic Acids Res 37, 1–13.

Huysmans, M., Lema, A.S., Coll, N.S., and Nowack, M.K. (2017). Dying two deaths - programmed cell death regulation in development and disease. Curr Opin Plant Biol 35, 37–44.

Iida, H., Mahonen, A.P., Jurgens, G., and Takada, S. (2023). Epidermal injury-induced derepression of key regulator ATML1 in newly exposed cells elicits epidermis regeneration. Nat Commun 14, 1031.

Iida, H., Yoshida, A., and Takada, S. (2019). ATML1 activity is restricted to the outermost cells of the embryo through post-transcriptional repressions. Development 146.

Ikeuchi, M., Favero, D.S., Sakamoto, Y., Iwase, A., Coleman, D., Rymen, B., and Sugimoto, K. (2019). Molecular Mechanisms of Plant Regeneration. Annu Rev Plant Biol 70, 377–406.

Ikeuchi, M., Iwase, A., Rymen, B., Lambolez, A., Kojima, M., Takebayashi, Y., Heyman, J., Watanabe, S., Seo, M., De Veylder, L., et al. (2017). Wounding Triggers Callus Formation via Dynamic Hormonal and Transcriptional Changes. Plant Physiol 175, 1158–1174.

Iwase, A., Kondo, Y., Laohavisit, A., Takebayashi, A., Ikeuchi, M., Matsuoka, K., Asahina, M., Mitsuda, N., Shirasu, K., Fukuda, H., et al. (2021). WIND transcription factors orchestrate wound-induced callus formation, vascular reconnection and defense response in Arabidopsis. New Phytol 232, 734–752.

Kawano, T. (2003). Roles of the reactive oxygen species-generating peroxidase reactions in plant defense and growth induction. Plant Cell Rep 21, 829–837.

Kim, D., Paggi, J.M., Park, C., Bennett, C., and Salzberg, S.L. (2019). Graph-based genome alignment and genotyping with HISAT2 and HISAT-genotype. Nat Biotechnol 37, 907–915.

Kim, H., Choi, D., and Suh, M.C. (2017). Cuticle ultrastructure, cuticular lipid composition, and gene expression in hypoxia-stressed Arabidopsis stems and leaves. Plant Cell Rep 36, 815–827.

Kim, M., Hyeon, D.Y., Kim, K., Hwang, D., and Lee, Y. (2024). Phytohormonal regulation determines the organization pattern of shoot aerenchyma in greater duckweed (Spirodela polyrhiza). Plant Physiol.

Kolde, R. (2019). Pheatmap: pretty heatmaps. R package version 1, 726.

Koo, A.J., and Howe, G.A. (2009). The wound hormone jasmonate. Phytochemistry 70, 1571–1580.

Kunst, L., and Samuels, A.L. (2003). Biosynthesis and secretion of plant cuticular wax. Prog Lipid Res 42, 51–80.

Langfelder, P., and Horvath, S. (2008). WGCNA: an R package for weighted correlation network analysis. BMC Bioinformatics 9, 559.

Lee, E.J., Kim, K.Y., Zhang, J., Yamaoka, Y., Gao, P., Kim, H., Hwang, J.U., Suh, M.C., Kang, B., and Lee, Y. (2021). Arabidopsis seedling establishment under waterlogging requires ABCG5-mediated for mation of a dense cuticle layer. New Phytol 229, 156–172.

Lee, Y., Rubio, M.C., Alassimone, J., and Geldner, N. (2013). A mechanism for localized lignin deposition in the endodermis. Cell 153, 402–412.

Lee, Y., Yoon, T.H., Lee, J., Jeon, S.Y., Lee, J.H., Lee, M.K., Chen, H., Yun, J., Oh, S.Y., Wen, X., et al. (2018). A Lignin Molecular Brace Controls Precision Processing of Cell Walls Critical for Surface Integrity in Arabidopsis. Cell 173, 1468–1480 e1469.

Leon, J., Rojo, E., and Sanchez-Serrano, J.J. (2001). Wound signalling in plants. J Exp Bot 52, 1–9.

Li, Y., Beisson, F., Koo, A.J., Molina, I., Pollard, M., and Ohlrogge, J. (2007). Identification of acyltrans ferases required for cutin biosynthesis and production of cutin with suberin-like monomers. Proc Natl Acad Sci U S A 104, 18339–18344.

Liu, J., Zhang, Z., Yu, Z., Liang, Y., Li, X., and Ren, L. (2018). Experimental study and numerical simulation on the structural and mechanical properties of Typha leaves through multimodal microscopy approaches. Micron 104, 37–44.

Livak, K.J., and Schmittgen, T.D. (2001). Analysis of relative gene expression data using real-time quantitative PCR and the 2(-Delta Delta C(T)) Method. Methods 25, 402–408.

Lu, P., Porat, R., Nadeau, J.A., and O’Neill, S.D. (1996). Identification of a meristem L1 layer-specific gene in Arabidopsis that is expressed during embryonic pattern formation and defines a new class of homeobox genes. The Plant Cell 8, 2155–2168.

Lu, S., Song, T., Kosma, D.K., Parsons, E.P., Rowland, O., and Jenks, M.A. (2009). Arabidopsis CER 8 encodes LONG-CHAIN ACYL-COA SYNTHETASE 1 (LACS1) that has overlapping functions with LACS2 in plant wax and cutin synthesis. Plant J 59, 553–564.

Lulai, E.C., and Neubauer, J.D. (2014). Wound-induced suberization genes are differentially expressed, spatially and temporally, during closing layer and wound periderm formation. Postharvest Biol Tec 90, 24–33.

Lulai, E.C., Suttle, J.C., and Pederson, S.M. (2008). Regulatory involvement of abscisic acid in potato tuber wound-healing. Journal of Experimental Botany 59, 1175–1186.

Lup, S.D., Tian, X., Xu, J., and Perez-Perez, J.M. (2016). Wound signaling of regenerative cell reprogramming. Plant Sci 250, 178–187.

Marhava, P., Hoermayer, L., Yoshida, S., Marhavy, P., Benkova, E., and Friml, J. (2019). Re-activation of Stem Cell Pathways for Pattern Restoration in Plant Wound Healing. Cell 177, 957–969 e913.

Martinez-Trujillo, M., Limones-Briones, V., Cabrera-Ponce, J.L., and Herrera-Estrella, L. (2004). Improving transformation efficiency of by modifying the floral dip method. Plant Mol Biol Rep 22, 63–70.

Matsuoka, K., Sato, R., Matsukura, Y., Kawajiri, Y., Iino, H., Nozawa, N., Shibata, K., Kondo, Y., Satoh, S., and Asahina, M. (2021). Wound-inducible ANAC071 and ANAC096 transcription factors promote cambial cell formation in incised Arabidopsis flowering stems. Commun Biol 4, 369.

McConn, M., Creelman, R.A., Bell, E., Mullet, J.E., and Browse, J. (1997). Jasmonate is essential for insect defense in Arabidopsis. Proceedings of the National Academy of Sciences 94, 5473–5477.

McFarlane, H.E., Shin, J.J., Bird, D.A., and Samuels, A.L. (2010). Arabidopsis ABCG transporters, which are required for export of diverse cuticular lipids, dimerize in different combinations. Plant Cell 22, 3066–3075.

Miller, G., Schlauch, K., Tam, R., Cortes, D., Torres, M.A., Shulaev, V., Dangl, J.L., and Mittler, R. (2009). The plant NADPH oxidase RBOHD mediates rapid systemic signaling in response to diverse stimuli. Sci Signal 2, ra45.

Mittler, R., Zandalinas, S.I., Fichman, Y., and Van Breusegem, F. (2022). Reactive oxygen species signalling in plant stress responses. Nat Rev Mol Cell Biol 23, 663–679.

Moon, G.J., Peterson, C.A., and Peterson, R.L. (1984). Structural, Chemical, and Permeability Changes Following Wounding in Onion Roots. Can J Bot 62, 2253-&.

Mousavi, S.A., Chauvin, A., Pascaud, F., Kellenberger, S., and Farmer, E.E. (2013). GLUTAMATE RECEPTOR-LIKE genes mediate leaf-to-leaf wound signalling. Nature 500, 422–426.

Nawrath, C. (2002). The biopolymers cutin and suberin. Arabidopsis Book 1, e0021.

Ogawa, E., Yamada, Y., Sezaki, N., Kosaka, S., Kondo, H., Kamata, N., Abe, M., Komeda, Y., and Takahashi, T. (2015). ATML1 and PDF2 Play a Redundant and Essential Role in Arabidopsis Embryo Development. Plant Cell Physiol 56, 1183–1192.

Ordinola-Zapata, R., Martins, J.N.R., Bramante, C.M., Villas-Boas, M.H., Duarte, M.H., and Versiani, M.A. (2017). Morphological evaluation of maxillary second molars with fused roots: a micro-CT study. Int Endod J 50, 1192–1200.

Orman-Ligeza, B., Parizot, B., de Rycke, R., Fernandez, A., Himschoot, E., Van Breusegem, F., Bennett, M.J., Perilleux, C., Beeckman, T., and Draye, X. (2016). RBOH-mediated ROS production facilitates lateral root emergence in Arabidopsis. Development 143, 3328–3339.

Park, C.S., Go, Y.S., and Suh, M.C. (2016). Cuticular wax biosynthesis is positively regulated by WRINKLED4, an AP2/ERF-type transcription factor, in Arabidopsis stems. Plant J 88, 257–270.

Park, J.H., Halitschke, R., Kim, H.B., Baldwin, I.T., Feldmann, K.A., and Feyereisen, R. (2002). A knock-out mutation in allene oxide synthase results in male sterility and defective wound signal transduction in Arabidopsis due to a block in jasmonic acid biosynthesis. Plant J 31, 1–12.

Pertea, M., Pertea, G.M., Antonescu, C.M., Chang, T.C., Mendell, J.T., and Salzberg, S.L. (2015). StringTie enables improved reconstruction of a transcriptome from RNA-seq reads. Nat Biotechnol 33, 29 0–295.

Peterson, K.M., Shyu, C., Burr, C.A., Horst, R.J., Kanaoka, M.M., Omae, M., Sato, Y., and Torii, K.U. ( 2013). Arabidopsis homeodomain-leucine zipper IV proteins promote stomatal development and ectopically induce stomata beyond the epidermis. Development 140, 1924–1935.

Pyke, K. (2009). Plastid biology (Cambridge University Press).

Qin, B.X., Tang, D., Huang, J., Li, M., Wu, X.R., Lu, L.L., Wang, K.J., Yu, H.X., Chen, J.M., Gu, M.H., et al. (2011). Rice OsGL1-1 is involved in leaf cuticular wax and cuticle membrane. Mol Plant 4, 985–995.

Rantong, G., Evans, R., and Gunawardena, A.H. (2015). Lace plant ethylene receptors, AmERS1a and AmERS1c, regulate ethylene-induced programmed cell death during leaf morphogenesis. Plant Mol Biol 89, 215–227.

Rittinger, P.A., Biggs, A.R., and Peirson, D.R. (1987). Histochemistry of Lignin and Suberin Deposition in Boundary-Layers Formed after Wounding in Various Plant-Species and Organs. Can J Bot 65, 18 86–1892.

Robinson, M.D., McCarthy, D.J., and Smyth, G.K. (2010). edgeR: a Bioconductor package for differential expression analysis of digital gene expression data. Bioinformatics 26, 139–140.

Rojas-Murcia, N., Hematy, K., Lee, Y., Emonet, A., Ursache, R., Fujita, S., De Bellis, D., and Geldner, N. (2020). High-order mutants reveal an essential requirement for peroxidases but not laccases in Casparian strip lignification. Proc Natl Acad Sci U S A 117, 29166–29177.

Sabba, R.P., and Lulai, E.C. (2002). Histological analysis of the maturation of native and wound periderm in potato (Solanum tuberosum L.) Tuber. Ann Bot 90, 1–10.

Sagi, M., and Fluhr, R. (2006). Production of reactive oxygen species by plant NADPH oxidases. Plant Physiol 141, 336–340.

San-Bento, R., Farcot, E., Galletti, R., Creff, A., and Ingram, G. (2014). Epidermal identity is maintained by cell-cell communication via a universally active feedback loop in Arabidopsis thaliana. Plant J 77, 46–58.

Savatin, D.V., Gramegna, G., Modesti, V., and Cervone, F. (2014). Wounding in the plant tissue: the defense of a dangerous passage. Front Plant Sci 5, 470.

Schmieder, R., and Edwards, R. (2011). Quality control and preprocessing of metagenomic datasets. Bioinformatics 27, 863–864.

Schnurr, J., Shockey, J., and Browse, J. (2004). The acyl-CoA synthetase encoded by LACS2 is essential for normal cuticle development in Arabidopsis. The Plant Cell 16, 629–642.

Schreiber, L., Franke, R., and Hartmann, K. (2005). Wax and suberin development of native and wound periderm of potato (Solanum tuberosum L.) and its relation to peridermal transpiration. Planta 220, 520–530.

Serra, O., and Geldner, N. (2022). The making of suberin. New Phytol 235, 848–866.

Serra, O., Mahonen, A.P., Hetherington, A.J., and Ragni, L. (2022). The Making of Plant Armor: The Periderm. Annu Rev Plant Biol 73, 405–432.

Sessions, A., Weigel, D., and Yanofsky, M.F. (1999). The Arabidopsis thaliana MERISTEM LAYER 1 promoter specifies epidermal expression in meristems and young primordia. The Plant Journal 20, 259–263.

Straus, M.R., Rietz, S., Ver Loren van Themaat, E., Bartsch, M., and Parker, J.E. (2010). Salicylic acid antagonism of EDS1-driven cell death is important for immune and oxidative stress responses in Arabidopsis. Plant J 62, 628–640.

Suh, M.C., Samuels, A.L., Jetter, R., Kunst, L., Pollard, M., Ohlrogge, J., and Beisson, F. (2005). Cuticular lipid composition, surface structure, and gene expression in Arabidopsis stem epidermis. Plant Physiol 139, 1649–1665.

Takada, S. (2013). Post-embryonic induction of ATML1-SRDX alters the morphology of seedlings. PLoS One 8, e79312.

Takada, S., Takada, N., and Yoshida, A. (2013). ATML1 promotes epidermal cell differentiation in Arabidopsis shoots. Development 140, 1919–1923.

Takahashi, K., Shimada, T., Kondo, M., Tamai, A., Mori, M., Nishimura, M., and Hara-Nishimura, I. (2010). Ectopic Expression of an Esterase, Which is a Candidate for the Unidentified Plant Cutinase, Causes Cuticular Defects in Arabidopsis thaliana. Plant Cell Physiol 51, 123–131.

Tao, X.Y., Mao, L.C., Li, J.Y., Chen, J.X., Lu, W.J., and Huang, S. (2016). Abscisic acid mediates wound-healing in harvested tomato fruit. Postharvest Biol Tec 118, 128–133.

Torres, M.A., Dangl, J.L., and Jones, J.D. (2002). Arabidopsis gp91phox homologues AtrbohD and AtrbohF are required for accumulation of reactive oxygen intermediates in the plant defense response. Proc Natl Acad Sci U S A 99, 517–522.

Tripathi, A.K., Pareek, A., and Singla-Pareek, S.L. (2017). TUNEL Assay to Assess Extent of DNA Fragmentation and Programmed Cell Death in Root Cells under Various Stress Conditions. Bio Protoc 7, e2502.

Tsuwamoto, R., Fukuoka, H., and Takahata, Y. (2008). GASSHO1 and GASSHO2 encoding a putative leucine-rich repeat transmembrane-type receptor kinase are essential for the normal development of the epidermal surface in Arabidopsis embryos. Plant J 54, 30–42.

Ursache, R., Andersen, T.G., Marhavy, P., and Geldner, N. (2018). A protocol for combining fluorescent proteins with histological stains for diverse cell wall components. Plant J 93, 399–412.

Völz, R., Heydlauff, J., Ripper, D., von Lyncker, L., and Groß-Hardt, R. (2013). Ethylene signaling is required for synergid degeneration and the establishment of a pollen tube block. Developmental Cell 25, 310–316.

Wang, H., Li, Y., Pan, J., Lou, D., Hu, Y., and Yu, D. (2017). The bHLH Transcription Factors MYC2, MYC3, and MYC4 Are Required for Jasmonate-Mediated Inhibition of Flowering in Arabidopsis. Mol Plant 10, 1461–1464.

Yang, W., Zhai, H., Wu, F., Deng, L., Chao, Y., Meng, X., Chen, Q., Liu, C., Bie, X., Sun, C., et al. (2024). Peptide REF1 is a local wound signal promoting plant regeneration. Cell 187, 3024–3038 e3014.

Zhang, G., Zhao, F., Chen, L., Pan, Y., Sun, L., Bao, N., Zhang, T., Cui, C.X., Qiu, Z., Zhang, Y., et al. (2019). Jasmonate-mediated wound signalling promotes plant regeneration. Nat Plants 5, 491–497.

Zhou, W., Lozano-Torres, J.L., Blilou, I., Zhang, X., Zhai, Q., Smant, G., Li, C., and Scheres, B. (2019). A Jasmonate Signaling Network Activates Root Stem Cells and Promotes Regeneration. Cell 177, 942–956 e914.

